# Unique, shared, and dominant brain activation in Visual Word Form Area and Lateral Occipital Complex during reading and picture naming

**DOI:** 10.1101/2021.09.20.461111

**Authors:** Josh Neudorf, Layla Gould, Marla J. S. Mickleborough, Chelsea Ekstrand, Ron Borowsky

## Abstract

Identifying printed words and pictures concurrently is ubiquitous in daily tasks, and so it is important to consider the extent to which reading words and naming pictures may share a cognitive-neurophysiological functional architecture. Two functional magnetic resonance imaging (fMRI) experiments examined whether reading along the left ventral occipitotemporal region (vOT; often referred to as a visual word form area, VWFA) has activation that is overlapping with referent pictures (i.e., both conditions significant and *shared*, or with one significantly more *dominant*) or *unique* (i.e., one condition significant, the other not), and whether picture naming along the right lateral occipital complex (LOC) has overlapping or *unique* activation relative to referent words. Experiment 1 used familiar regular and exception words (to force lexical reading) and their corresponding pictures in separate naming blocks, and showed *dominant* activation for pictures in the LOC, and *shared* activation in the VWFA for exception words and their corresponding pictures (regular words did not elicit significant VWFA activation). Experiment 2 controlled for visual complexity by superimposing the words and pictures and instructing participants to either name the word or the picture, and showed primarily *shared* activation in the VWFA and LOC regions for both word reading and picture naming, with some *dominant* activation for pictures in the LOC. Overall, these results highlight the importance of including exception words to force lexical reading when comparing to picture naming, and the significant *shared* activation in VWFA and LOC serves to challenge specialized models of reading or picture naming.

## Introduction

The co-appearance of words and pictures in everyday materials affect how we perceive, interpret, and understand information. Furthermore, words and pictures are often presented in conjunction to convey additional information over and above what either modality could convey if presented alone, such as when reading textbooks, manuals, magazines, graphic novels, and movie subtitles, playing video games, or viewing internet news, advertisements and memes. A growing number of studies have examined the possible neurobiological interactions between these modalities. Although there is some consensus that the processes underlying word reading and picture naming tasks both activate the ventral visual processing stream, there remains debate regarding whether there are particular subregions of the ventral stream that are more specialized for either word reading or picture naming or whether the underlying processes activate shared regions. Some researchers have described the visual system as being attuned to the requirements of each modality’s processes (i.e., functional specialization from a neuroanatomical perspective, e.g., Cohen et al., 2000, or from a neural network perspective, e.g., Masson & Borowsky, 1998), while others take the view that the visual system operates in a more domain-general fashion (e.g., Kherif et al., 2011; Price & Devlin, 2003, 2004).

Research evidence has been used to support theories about the specialization of brain regions in word reading processes. Cohen and colleagues have shown that words activate a specific region of the left mid-fusiform gyrus/occipitotemporal cortex, which they propose specializes in word form identification and abstract representations of visual words, and has led to that region being labeled as the ‘visual word form area’ (VWFA; Cohen et al., 2000; Dehaene et al., 2001, 2002; Dehaene & Cohen, 2011; see also Fiez & Petersen, 1998; Hart et al., 2000; Leff et al., 2001). In support of this description, Cohen et al. (2002) showed that the VWFA is a critical lesion site for pure alexia, as damage to this region may lead to a unimodal deficit in word reading (e.g., Beversdorf et al., 1997; Damasio & Damasio, 1983; Leff et al., 2001). It has also been shown at an individual participant level that passive viewing of English words produces more activation than line drawings (which did not correspond to the words), and more than words with unfamiliar characters, in a region of the left ventral occipital cortex in a majority of participants (Baker et al., 2007). There has also been some recent work involving word/nonword discrimination in VWFA, resting state functional and structural connectivity between VWFA and Wernicke’s area including semantic processing, and direct stimulation of VWFA in temporal lobe epilepsy patients that has revitalized this hypothesis about specialization for word reading in this region (Behrmann & Plaut, 2020; Stevens et al., 2017; Woolnough et al., 2020).

Other studies have shown that the VWFA is also a critical region involved in processing alternative types of visual stimuli. For example, it has been shown that nonwords have greater activation in the VWFA region than real words (Brunswick et al., 1999; Fujimaki et al., 1999; Tagamets et al., 2000; Xu, 2001). However, this finding could reflect a form of frequency effect, whereby lower frequency letter strings tend to produce greater activation than higher frequency letter strings in linguistically-relevant regions (e.g., Cummine et al., 2010). It has also been shown that processes involved in picture related tasks (e.g., semantic processing, naming, viewing, one-back memory task) can produce greater activation in regions of the VWFA than words (Chee et al., 2000; Moore & Price, 1999; Sevostianov et al., 2002; Vandenberghe et al., 1996; see also Price & Devlin, 2003, 2004; for a review see Price, 2012). Moore and Price (1999) also found regions of the VWFA were sensitive to meaningful stimuli more than control stimuli regardless of task (name or view) and regardless of stimuli (word or picture), and that regions of the VWFA were more sensitive to naming than viewing regardless of stimuli (word or picture). Neuropsychological reports of word processing have also found that the VWFA is more sensitive to naming than to non-verbal lexical tasks (e.g., Hillis et al., 2005). In contrast, some research using tasks such as a one-back task have produced activation for words greater than scrambled controls in a region of the VWFA in which no activation was observed for pictures greater than scrambled controls (Szwed et al., 2011). Another example of how task demands can alter differences between words and pictures in the VWFA includes comparisons between color decision and object categorization, whereby words produced more activation than pictures during the color decision task but not during the object categorization task (Starrfelt & Gerlach, 2007). A more recent study with a large adult sample with varying degrees of literacy showed that before literacy faces primarily produced activation in RH fusiform face area (FFA) and in LH VWFA, and that after literacy words also produced activation in LH VWFA, but that faces continued to activate in this region without suffering from competition with word-related activation (Hervais-Adelman et al., 2019). Findings of shared (or greater) activation for pictures compared to words in this region challenges the notion that this region specializes exclusively in visual word form. Lastly, there have also been many cases of patients with pure alexia that were reported to also have difficulties with picture related tasks and color naming (Behrmann et al., 1998; Damasio & Damasio, 1983; De Renzi et al., 1987; Geschwind, 1965).

In contrast, a primarily right hemisphere ventral region has been shown to respond to visual objects more so than other stimuli. A seminal study by Malach et al. (1995) showed that the lateral bank of the fusiform gyrus extending ventrally and dorsally, which was termed the lateral occipital complex (LOC), responded more strongly when subjects viewed photographs of objects than when they viewed visual textures. A similar result was demonstrated in a study by Kanwisher et al. (1996), in which they found that drawings elicited stronger responses in the LOC than scrambled counterparts. Furthermore, it has been suggested that the LOC is an object-selective region, as it shows a number of responsive properties that characterize an effective object recognition system (Doniger et al., 2000; Grill-Spector et al., 1999; Kourtzi, 2001; Murtha et al., 1999; Snow et al., 2011). Results from lesion studies provide evidence that the LOC may be essential for object recognition in that damage to this region often results in a variety of object recognition deficits (Damasio et al., 1990; Farah et al., 1989; Farah et al., 1991, 1995; Feinberg et al., 1994; Goodale et al., 1991; Moscovitch et al., 1997). Although the LOC has been strongly implicated in object related tasks, it is important to investigate whether this region is also involved in word reading (see also Nestor et al., 2013).

Although reading and picture naming are similar tasks in that they both involve identifying visual stimuli and generating verbal responses, there is also a key difference between the tasks. That is, word reading can involve lexical processing (i.e., sight word reading through an orthographic lexical system) and sublexical processing (i.e., grapheme to phoneme conversion), whereas picture naming involves recognizing the object from memory (see Fig 1). Moreover, in word reading tasks, regular words (REG; which follow regular spelling-to-sound mappings; e.g., coin,) can be processed either lexically or sublexically, whereas exception words (EXC; which follow irregular spelling-to-sound mappings; e.g., comb) must be processed lexically in order to be pronounced correctly. Thus, previous studies that have examined whether words and pictures share common functional regions may be qualified by the fact that they did not control for the degree of lexical reliance in reading through using EXC words, including some of our prior research (Borowsky et al., 2005, see also Cummine et al., 2013 for additional approaches for manipulating lexical reliance). Furthermore, our previous research comparing picture naming to EXC word reading (Borowsky et al., 2007) did not utilize matched picture and word stimuli, which we employ in the present research in order to provide the best-controlled comparison of lexical reading and picture naming. Experiments 1 and 2 involved using fMRI to examine activation for matched picture and word stimuli in word reading and picture naming, including stimuli that optimally activate the ventral-lexical stream (i.e., EXC words), thus controlling for the degree of lexical-based reading and allowing for an assessment of shared versus unique activation loci relative to picture versions of the same referents. We also examined the same regions using familiar REG words and their corresponding pictures.

**Fig 1.**
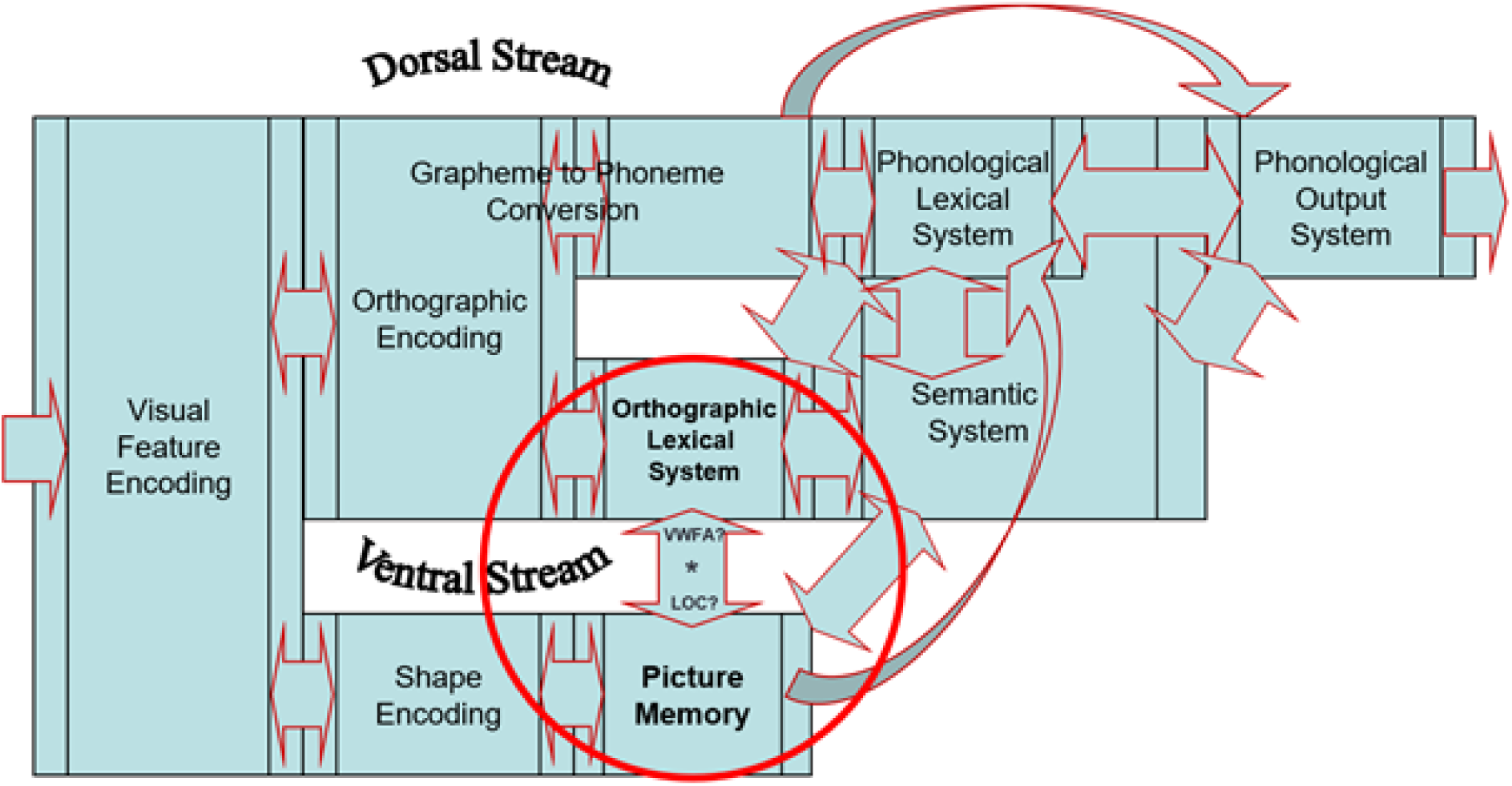
Dual route model of reading (top) adapted from Coltheart et al. (2001); Cummine et al. (2010, 2013); and Gould et al. (2012), merged with a basic model of picture naming (bottom) adapted from Paivio (1990), see Johnson et al. (1996) for a review.

One important aspect to consider when comparing words and pictures is the differences in visual complexity (Szwed et al., 2011). Although picture and word stimuli can be adequately matched on several variables (e.g., luminance, shape, and visual angle), other disparities between pictures and words cannot be as easily controlled (e.g., visual detail, spatial frequency, and color). To address this, Experiment 1 compared the activation for word and picture naming in isolation and Experiment 2 superimposed the stimuli in order to equate for overall visual complexity. The latter paradigm was a task in which a word was superimposed on a picture and participants were instructed to name either the word or the picture (de Zubicaray et al., 2002; Glaser & Düngelhoff, 1984; Posnansky & Rayner, 1977; Rosinski et al., 1975; Schriefers et al., 1990). Thus, in Experiment 2 the stimuli were perfectly matched not only for phonology (i.e., pronunciation) and semantics (i.e., meaning), but also for visual complexity as the same stimulus was used for both the word and picture naming conditions. Importantly, a comparison using EXC words, which are optimal for activating lexical representations, and their corresponding pictures, has not yet been investigated.

Another important consideration is to clearly distinguish *unique* activation from overlapping activation that is either *shared*, or *dominant* for one of the processes. *Unique* activation can be defined whereby one of the conditions is significant, but the other is not, and the difference in activation between the conditions is significant. Overlapping activation can be of two types: 1) *shared*, whereby both conditions are each significant but do not significantly differ (e.g., Borowsky et al., 2005, 2006, 2007; see also Price & Friston, 1997), or 2) *dominant*, whereby both conditions are each significant but one is significantly greater than the other. If words and pictures share a common functional architecture, there should be some form of overlapping activation for words and pictures. Alternatively, if words and pictures activate *unique* functional regions, there should be *unique* activation for words in the VWFA region, and *unique* activation for pictures in the LOC.

As previously mentioned, the degree to which words and pictures may co-activate or uniquely activate the VWFA and LOC is contentious, and previous research has not examined optimal stimuli such as EXC words and their corresponding matching pictures. The present experiments examined whether naming of words and pictures activate the same regions, including high frequency EXC words and their corresponding pictures, which should be the optimal stimuli for activating the ventral-lexical stream, as well as high frequency REG words and pictures. Finally, overt naming was used so as to maintain a high degree of ecological validity, and we have previously demonstrated the effectiveness of a sparse-coding fMRI paradigm for allowing participants to make spoken responses (Borowsky et al., 2006, 2007; Cummine et al., 2010, 2013; Gould et al., 2017, 2018). Although the basic design for each experiment is a 2 (Word type: EXC vs. REG) X 2 (Stimulus type: Word vs. Picture) factorial design, in order to fully assess the degree of unique versus overlapping (shared or dominant) activation in the regions of interest, we focus on specific matched contrasts (e.g., EXC words vs. EXC pictures in the VWFA). The hypothesis that the VWFA specializes in processing word stimuli would be supported if significant unique or dominant activation for words is found in that region compared to pictures, whereas the hypothesis that the LOC specializes in processing picture stimuli would be supported if significant unique or dominant activation for pictures is found in that region compared to words. Given that EXC word stimuli are optimal for activating orthographic lexical representations, the strongest test of these hypotheses should result from comparing EXC word and picture stimuli.

## Experimental Procedures

### Experiment 1

#### Participants

Fifteen healthy participants with normal or corrected-to-normal vision and fluent English participated in the experiment. The sample of participants were 12 females and 3 males (median age = 26 years, aged between 20 and 31, all right-handed). The participants’ informed, written consent was obtained, and the experiment was performed in compliance with the relevant laws and institutional guidelines and was approved by the University of Saskatchewan Research Ethics Board.

#### Stimuli

Fifty monosyllabic words (25 EXC, 25 REG) and their corresponding pictures (acquired using the corresponding word as the search term in Google images) were used as critical stimuli. The use of pictures that corresponded to these EXC and REG words is a novel approach in the context of neuroimaging research of the VWFA and LOC that allows for the control of semantic and phonological differences between word and picture naming. These stimuli were matched on several of the characteristics available from the E-Lexicon Database (http://elexicon.wustl.edu/; Balota et al., 2007), and we verified that word type (REG vs. EXC) was not associated with log_10_ HAL word frequency (*r* = -.13, *p* = .36), bigram frequency by position (*r* = -.04, *p* = .77), length (*r =* -.09, *p* = .54), orthographic neighbors (*r* = -.03, *p* = .86), number of phonemes (*r* = -.07, *p* = .65), or semantic neighborhood density (Shaoul & Westbury, 2010; *r* = -.02, *p* = .91). These words are likely of high familiarity for participants, as their mean corpus-based word frequency (log_10_ HAL word frequency mean = 2.86) is relatively high. In order to obtain ratings for how well the picture names and words agree with one another, a separate group of 40 undergrad participants were asked to rate the stimuli as part of a separate study. For these ratings, a picture and its corresponding word were presented on the computer screen and participants were asked to “rate how well the picture matches the label” on a scale of one (very poorly) to five (very well). The reliability of these subjective ratings was estimated using the Spearman-Brown formula, which gave a coefficient of .91, indicating good reliability. This measure was not confounded with Word Type (i.e., REG versus EXC), as a point-biserial correlation was computed and found to be r = -.084, p = .52. This indicates that picture-orthography agreement was not significantly related to Word Type. The words subtended a horizontal visual angle of between 13 and 19 degrees, and a vertical angle of 4 degrees. The pictures subtended a horizontal visual angle of between 7 and 18 degrees, and a vertical visual angle of between 9 and 18 degrees.

#### Protocol

All imaging was conducted using a 3 Tesla Siemens Skyra scanner. Whole-brain anatomical scans were acquired using high resolution axial magnetization prepared rapid acquisition gradient echo (MPRAGE) sequence, consisting of 192 T1-weighted images of 1 mm thickness (no gap) with an in-plane resolution of 1 x 1 mm (field of view 256; TR = 1900 ms; TE = 2.1 ms). For the functional scans, T2*-weighted single-shot gradient-echo echo-planar imaging (EPI) scans were acquired using an interleaved ascending EPI sequence, consisting of 55 volumes of 25 axial slices of 4 mm thickness (1 mm gap) with an in-plane resolution of 2.7 x 2.7 mm (field of view = 250), using a flip angle of 90**°**. The top 2 coil sets (16 channels) of a 20-channel Siemens head-coil were used, with the bottom set for neck imaging (4 channels) turned off. Additional foam padding was used to reduce head motion. In order to obtain verbal behavioural data from the MRI, we used a sparse-sampling (gap paradigm) fMRI method that allows the participant to respond during a gap in image acquisition (TR = 3300 ms, with a 1650 ms gap of no image acquisition, TE = 30 ms). A within-subjects design was used, and participants responded vocally during the regular, periodic gap in the image acquisition that followed the offset of each volume of image acquisition, which allowed the participants to respond with no noise interference from the MRI. That is, a stimulus was presented at the offset of an image acquisition for 1650 ms, providing a silent gap for participants to name aloud the stimulus.

#### Procedure and Apparatus

Stimuli were presented to participants in the center of a screen using a PC running EPrime software (Psychology Software Tools, Inc., http://www.pstnet.com) through MRI compatible goggles (Cinemavision Inc., http://www.cinemavision.biz). Each participant was presented with two runs of words (containing five blocks of five stimuli each and five relaxation blocks), and the order of these runs were counterbalanced by word type. The order of stimulus presentation within blocks was randomized. Given the possible ambiguity in picture naming and the desire for keeping the error rates low and comparable to the words, participants were presented with two runs of pictures, which were also counterbalanced by word type, subsequent to the word runs (see Fig 2). By naming the words first the participants were effectively primed to name the corresponding picture with the same verbal response. Participants were instructed to name aloud the stimulus as quickly and accurately as possible. For each of the four runs (REG Word, EXC Word, REG Picture, EXC Picture), 25 stimuli were presented in a random order in blocks of five stimuli, with each block followed by a relaxation block of equivalent duration (16.5 seconds for each block of stimuli or relaxation; relaxation blocks involved staring at the fixation cross, see Fig 3). One stimulus was presented for each TR. The leading edge (10 μ ec) of the fiber-optic signal that is emitted by the MRI at the beginning of each acquisition volume was detected by a Siemens fMRI trigger converter and passed to the Eprime PC via the serial port. In this way, perfect continuous synchronization between the MRI and the experimental paradigm computer was obtained at each volume. In order to avoid head movement while speaking, the participants were instructed to speak with their mouth kept slightly open to minimize mouth and jaw movements and were encouraged to not swallow or lick their lips during the experimental trials.

**Fig 2.**
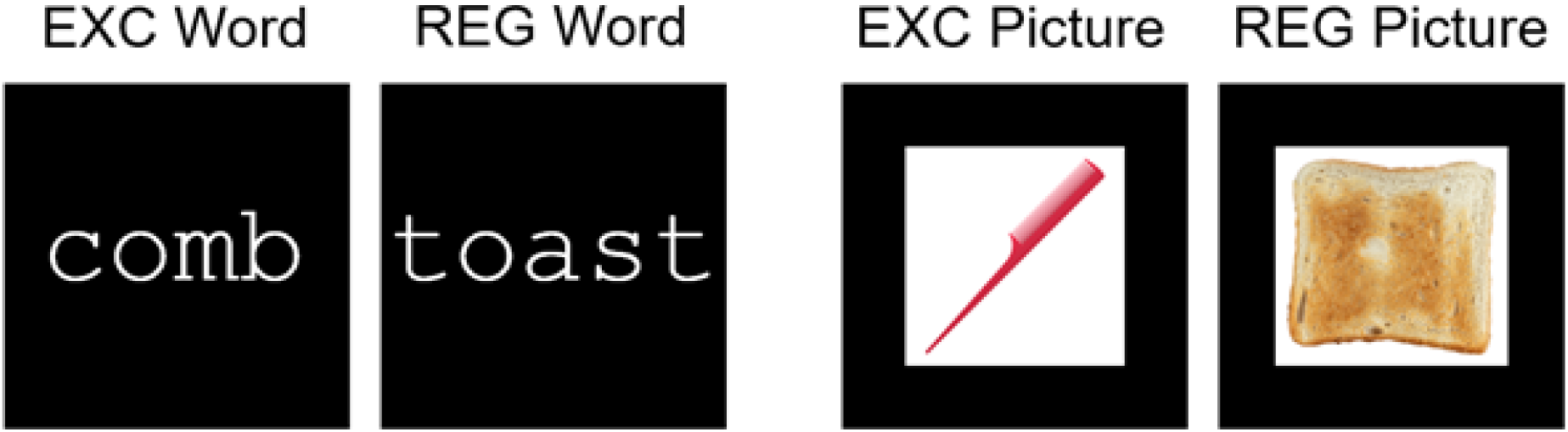
Stimuli examples for all four trial types of EXC words, REG words, EXC pictures, and REG pictures. These examples are for illustrative purposes only.

**Fig 3.**
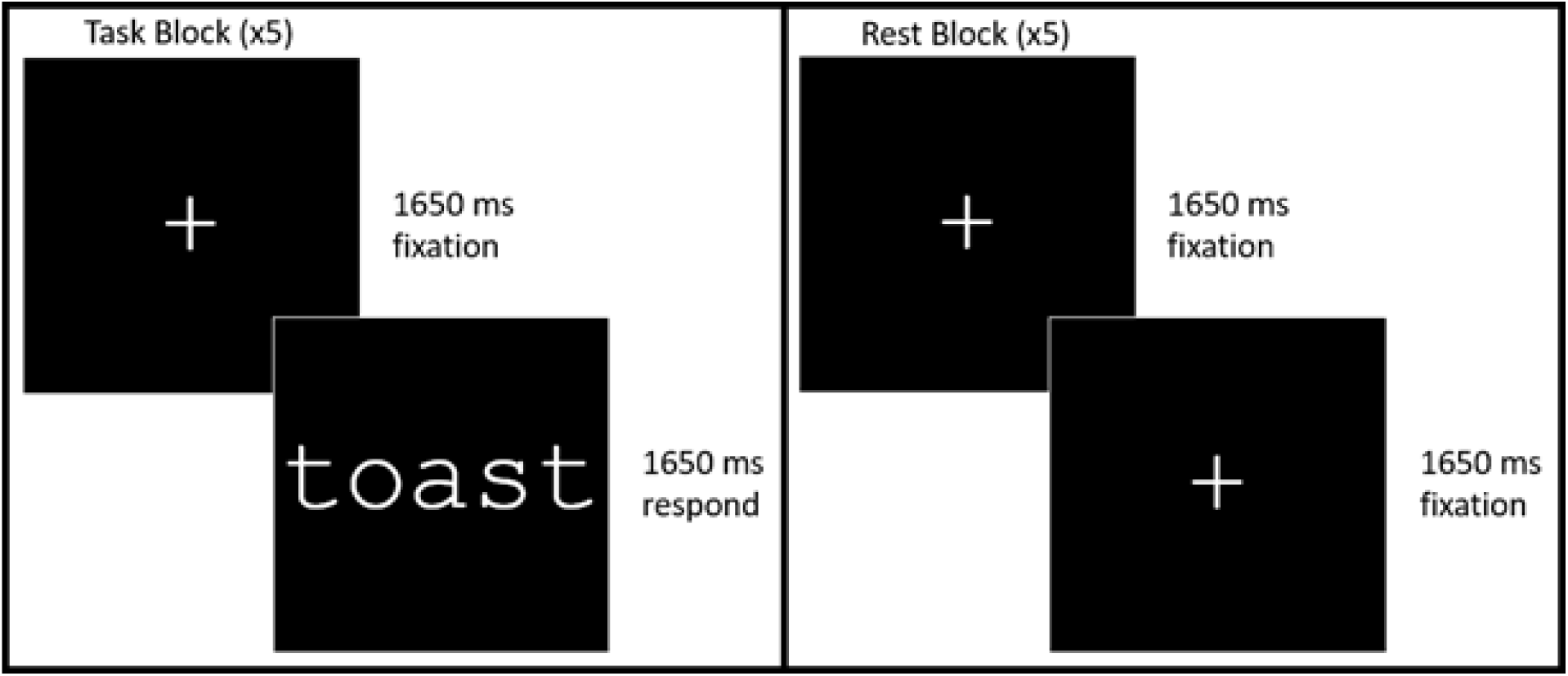
Experiment 1 trial progression example for REG word stimulus, demonstrating task and rest blocks.

Vocal responses were recorded at 96KHz, 24bit using an Olympus LS11 digital recorder. These recordings were analyzed using PRAAT software (Boersma & Weenink, 2009), and the waveforms and broadband spectrograms were used to localize vocalization reaction time (RT). The sparse-sampling (gap paradigm) fMRI method allowed the participants to make overt vocalizations in the absence of scanner noise during a gap in image acquisition, allowing RT data to be collected. The stimulus onset was synchronized with the last gradient sound before the gap. This provided an acoustic marker for the stimulus onset on the recording while the gap allowed for a clear recording of the participant’s vocal response.

#### FMRI Analysis

All preprocessing and statistical analyses for functional images were performed using FMRIB Software Library (FSL; Jenkinson et al., 2012). Functional images were preprocessed including slice scan time acquisition correction, 3D motion correction, spatial smoothing with a 5mm Full Width Half Maximum (FWHM) gaussian filter, and temporal filtering with a high-pass filter to filter frequencies lower than one complete block/rest cycle (10 TRs; period of 33 seconds). Functional volumes were then registered to anatomical brain images using FLS *flirt* with 7 degrees of freedom before being registered to standard MNI space with 12 degrees of freedom. Motion parameters were regressed as variables in the model to eliminate any artifacts from motion. The first-level analysis was conducted separately on time course data from all 4 runs (EXC Picture, EXC Word, REG Picture, REG Word). Each of these conditions was compared to the resting condition in order to produce the first-level activation map for each of the 4 runs/conditions. Contrasts were calculated in the second-level analysis at the subject level for EXC Picture > EXC Word, EXC Word > EXC Picture, REG Picture > REG Word, REG Word > REG Picture, and basic effects of EXC Picture, EXC Word, REG Picture, and REG Word. For the group-level analysis, a mean of the first-level (EXC Picture, EXC Word, REG Picture, REG Word) and second-level (EXC Picture > EXC Word, EXC Word > EXC Picture, REG Picture > REG Word, REG Word > REG Picture) contrasts was calculated. Additionally, a factorial group-level analysis was conducted on the first-level analysis outputs, which had a design of 2 (Word Type; EXC vs. REG) x 2 (Stimulus Type; Picture vs. Word), including an interaction term of Word Type x Stimulus Type. Statistics were calculated using FSL *randomise* with 5000 permutations, Threshold Free Cluster Enhancement (TFCE), and family-wise error correction at *p* < .01 (Nichols & Holmes, 2002; Smith & Nichols, 2009; Winkler et al., 2014).

An additional voxelwise individual participant-level analysis was conducted on a second-level Words > Pictures contrast to determine whether this contrast was significant in a region of interest (ROI) sphere of 10mm radius centered on the VWFA (MNI: *x* = -46, *y* = -53, *z* = -20) for a significant number of participants. Likewise this ROI localizer analysis was conducted on a second-level Pictures > Words contrast to determine whether this contrast was significant in a ROI sphere of 10mm radius centered on the LOC (MNI: *x* = 39, *y* = -75, *z* = -9) for a significant number of participants. The analysis was identical to the lower-level analysis used at the group level, but the Words > Pictures and Pictures > Words contrasts were examined voxelwise in the corresponding ROI for each individual. A pre-threshold mask for VWFA or LOC was applied in FSL *feat*, which then applied a multiple comparison correction based on the number of voxels in the mask and taking into account that neighboring voxels are not independent. A significance threshold for the number of participants showing the effect was chosen based on the binomial distribution, which requires at least 12 observations out of 15 to be significant at *p* = .05. The underlying assumption on which a ROI localizer approach is based is that there is a region associated with a cognitive process that is consistently activated across individuals, but that the precise location may vary to some small extent. If a significant number of individuals cannot be shown to have activation in the vicinity of a ROI then the logic of a localizer approach is not valid. Conversely, a group level GLM analysis does not make this assumption.

### Experiment 2

The same methods were used as in Experiment 1 with the following exceptions:

#### Participants

An additional 15 healthy participants with normal or corrected-to-normal vision and fluent English participated in Experiment 2. The sample of participants were 10 females and 5 males (median age = 27 years, aged between 21 and 46, all right-handed).

#### Stimuli

Participants viewed the superimposed word-picture stimuli, whereby a grey font was used so that the word would be visible on both dark and light backgrounds (see Fig 4; see Fig 5 for trial progression). The stimulus was either congruent or incongruent to the corresponding referent (50% congruent), and they were asked to either name the words (in the word run) or the pictures (in the picture run) aloud as quickly and accurately as possible. Incongruent trials were included to force participants to rely on the words in the word block and on the pictures in the picture block.

**Fig 4.**
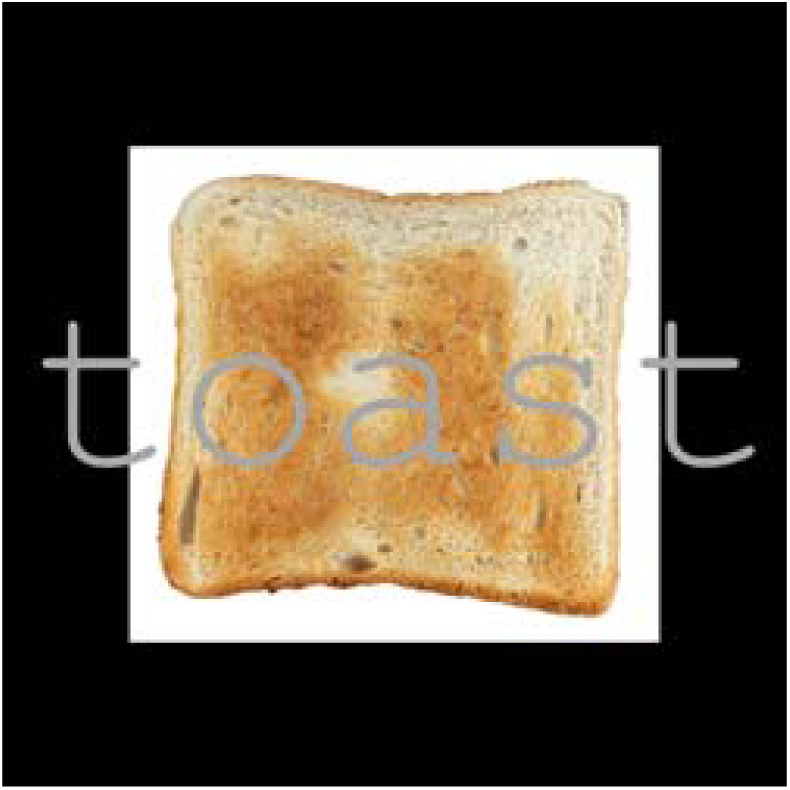
An example of a superimposed stimulus used where participants were instructed to name either the word or the picture. This stimulus is similar but not identical to the original image and is therefore for illustrative purposes only.

**Fig 5.**
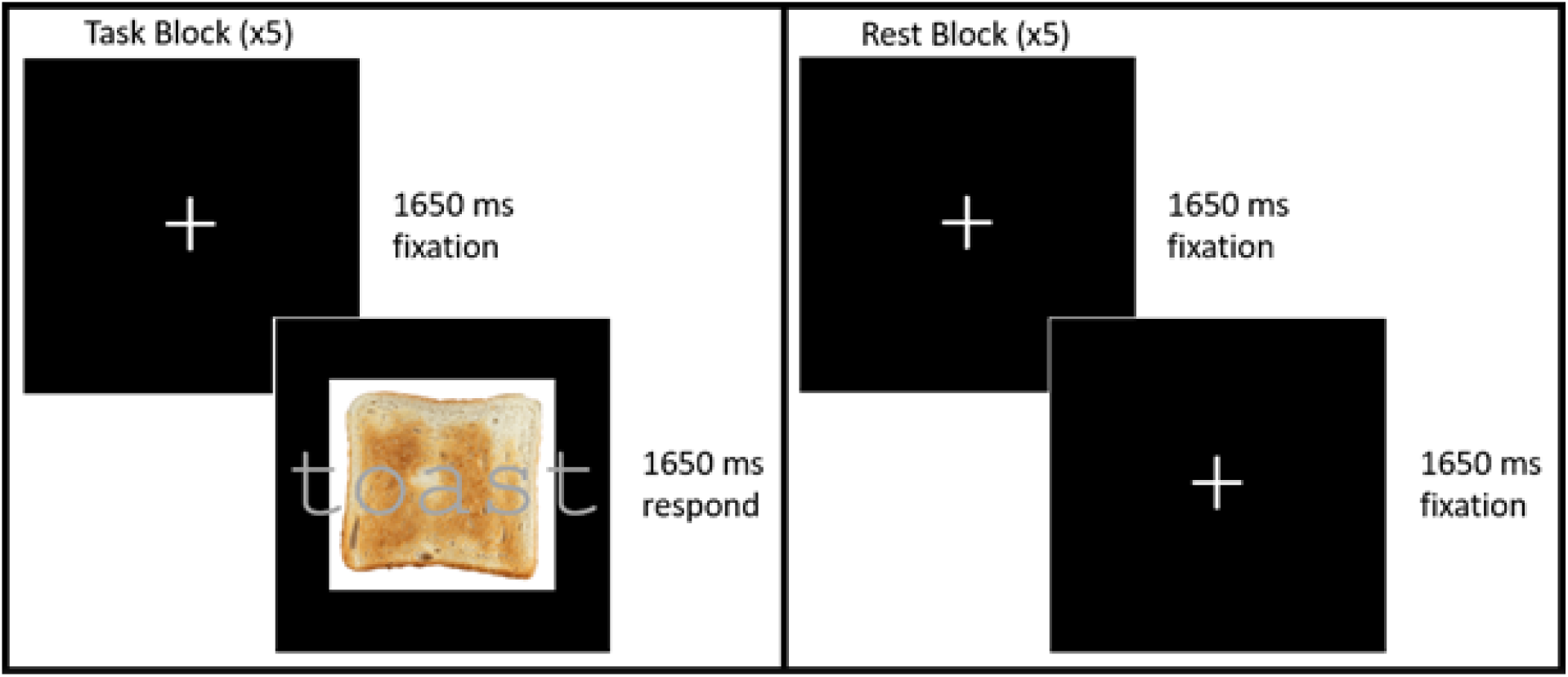
Experiment 2 trial progression example for REG word/picture stimulus, demonstrating task and rest blocks.

#### Procedure and Apparatus

In Experiment 2, the blocks of words and pictures were counterbalanced by Word Type and Stimulus Type to counteract any order effects.

## Results

### Experiment 1

#### FMRI Group Analyses

For both EXC and REG word types, the contrast of Pictures > Words produced a large amount of activation. We examined the higher involvement for picture naming compared to reading by investigating brain regions where the effect for picture was *unique* (pictures were significantly active compared to rest while words were not), *dominant* (both words compared to rest and pictures compared to rest were significantly active, but pictures were still more activated than words) or rather *shared* between pictures and words (both pictures and words were significantly activated and there was no statistically significant difference between them), for each word-type separately (see Fig 4). The *dominant*, *unique*, and *shared* effects were calculated as conjunction maps, whereby *dominant* was the conjunction [(Pictures ∩ Words) ∩ (Pictures > Words)], *unique* was the conjunction [(Pictures > Words) ∩ ∼Words], and *shared* was the conjunction [(Pictures ∩ Words) ∩ ∼(Pictures > Words)].

No regions of activation were found to be significant for Word > Pictures, neither in the basic contrasts of EXC Word > EXC Pictures and REG Word > REG Picture, nor in the factorial analysis. However, Picture > Word contrasts produced significant activation in the factorial analysis and when investigated at the EXC Pictures > EXC Word and REG Picture > REG Word level. These effects were investigated along with *shared* processing between the basic EXC Picture and EXC Word contrasts and also between the REG Picture and REG Word contrasts. Activation contrasts in the vicinity of the VWFA (Cohen et al., 2000; Talairach coordinates *x* = -43, *y* = -54, *z* = -12 converted to MNI coordinates *x* = -46, *y* = -53, *z* = -20 using the transform outlined in Lacadie et al., 2008) and LOC (Grill-Spector, 2003; Talairach coordinates *x* = 40, *y* = -74, *z* = -2 converted to MNI coordinates *x* = 39, *y* = -75, *z* = -9) are discussed.

The VWFA showed *shared* activation for EXC pictures and words (Fig 4a), while REG pictures showed *unique* activation in the VWFA (Fig 4b), counter to the idea that words should activate the VWFA more so than pictures. The *unique* REG picture activation in the VWFA is consistent with some past research showing greater picture than word activation in the VWFA (Chee et al., 2000; Moore & Price, 1999; Sevostianov et al., 2002; Vandenberghe et al., 1996; see also Price & Devlin, 2003, 2004; for a review see Price, 2012). One could interpret this finding as being due to greater complexity of stimuli for pictures than for words. However, we saw that for EXC stimuli the Pictures > Words contrast demonstrated significant *shared* activation in the VWFA, so it may be that lexically read EXC words optimally activate the VWFA compared to REG words, leading to activation on the same level as pictures, whereas REG word activation of the VWFA is less than EXC due to joint lexical and sublexical reading processes being recruited (note that a direct comparison of EXC and REG did not yield a significant contrast, so further research is needed to examine this theory).

Contrasts in the LOC demonstrate overlapping activation for words and pictures, but activation for pictures is *dominant* (significantly greater than activation for words; Figs 4c and 3d). These patterns of activation in the LOC suggest that although picture activation is *dominant*, reading of words also significantly utilizes this region.

Other notable areas of *shared* word and picture activation for both EXC and REG words include activation of semantic regions in the anterior temporal lobes (see Fig 4; see Patterson et al., 2007), activation of phonological regions in the inferior frontal gyri and insula (see Figs 4c and 4d; see Borowsky et al., 2006), and putamen activation in the basal ganglia (see Fig 4c; see Gould et al., 2017, 2018; see also review by Price, 2012, and Oberhuber et al., 2013).

#### FMRI Individual Analyses

To investigate whether the Words > Pictures contrast may have been significant at the individual participant level in the VWFA, a ROI localizer analysis was conducted with the VWFA as the center (MNI: x = -46, y = -53, z = -20) and a liberally sized radius of 10 mm (twice the approximate standard deviation of the VWFA location observed by Cohen et al., 2002). None of the 15 participants produced a significant Words > Pictures contrast within 10mm of the VWFA center. This finding is consistent with what was found at the group level, whereby no part of the VWFA produced a significant Words > Pictures contrast.

Using the same approach for the Pictures > Words contrast in the LOC, 14 participants produced a significant Pictures > Words contrast within 10mm of the LOC center, which exceeds the threshold of 12 out of 15 based on the binomial distribution (see Fig 5 and Table 1). This finding is also consistent with what was found at the group level, whereby a significant Pictures > Words contrast was observed in the LOC.

**Table 1.** Experiment 1 largest cluster coordinates (center of gravity; i.e. weighted average of the coordinates by activation intensity) of significant Pictures > Words activation within 10mm of LOC (MNI: x = 39, y = -75, z = -9). Fourteen of 15 participants had a significant contrast within a 10mm radius ROI centered on the LOC.

| Participant | X | Y | Z | Voxels | Max Z-value |
| --- | --- | --- | --- | --- | --- |
| 1 | 34.9 | -76.1 | -10.6 | 52 | 5.77 |
| 2 | 35.0 | -71.1 | -12.3 | 17 | 6.24 |
| 3 | 40.8 | -74.5 | -11.0 | 310 | 7.44 |
| 5 | 39.2 | -75.7 | -10.0 | 348 | 9.66 |
| 6 | 32.6 | -75.1 | -11.8 | 39 | 5.70 |
| 7 | 40.1 | -76.9 | -10.5 | 265 | 8.73 |
| 8 | 36.2 | -72.4 | -11.1 | 90 | 7.00 |
| 9 | 40.6 | -79.7 | -11.2 | 172 | 7.27 |
| 10 | 42.7 | -81.4 | -9.7 | 24 | 5.96 |
| 11 | 39.8 | -74.7 | -9.1 | 136 | 5.74 |
| 12 | 36.5 | -76.7 | -10.1 | 230 | 8.93 |
| 13 | 36.8 | -81.6 | -6.0 | 5 | 4.10 |
| 14 | 43.1 | -77.8 | -7.2 | 108 | 8.50 |
| 15 | 40.1 | -75.8 | -10.1 | 501 | 12.80 |

#### Reaction Time

The RT data were aggregated by participant as a function of Stimulus Type (word, picture) and Word Type (REG, EXC). Medians of the correctly named item RTs were submitted to a 2 × 2 general linear model ANOVA, with Stimulus Type and Word Type as repeated measures factors. Due to equipment failure (i.e., faulty cable connector) one participant’s data was lost. There was a significant main effect of Stimulus Type, *F*(1, 13) = 167.68, *MSe* = *3569.01*, *p* < .001, whereby words (*M* = 658.08 ms) are named faster than pictures (*M* = 864.84 ms), which is consistent with previous reports in the literature (e.g., Fraisse, 1969; Hennessey & Kirsner, 1999; Potter & Faulconer, 1975). There was no significant main effect of Word Type, *F*(1, 13) = .25, *MSe* = 720.83, *p* = .62, and a marginally significant effect of Stimulus Type x Word Type interaction, *F*(1, 13) = 4.73, *MSe* = 1349.41, *p* = .05, whereby REG words (*M* = 649.20 ms) were named faster than EXC words (*M* = 666.96 ms), while REG pictures (*M =* 877.31 ms) were named slower than EXC pictures (*M* = 852.36 ms), with a 95% confidence interval (CI) for all means of 20.49 ms, using the within-subjects Loftus & Masson method (Masson & Loftus, 2003). Based on the 95% CI, REG pictures were significantly slower than EXC pictures (consistent with other research from our lab, Neudorf et al., in prep.), but the difference between REG and EXC words was not significant.

#### Error Rate

There was a significant main effect of Stimulus Type, *F*(1, 13) = 191.71, *MSe* = 70.71, *p* < .001, whereby there were fewer errors in the word reading conditions (*M* = 2.72%) than the picture naming conditions (*M* = 33.84%, notably higher as any response other than the paired word was considered an error). There was no significant main effect of Word Type, *F*(1, 13) = 1.674, *MSe* = 29.963, *p* = .22, and no significant Stimulus Type x Word Type interaction, *F*(1, 13) =3.01, *MSe* = 51.75, *p* = .11 (REG Words, *M* = 2.00%; EXC Words, *M* = 3.44%; REG Pictures, *M* = 36.45%; EXC Pictures, *M* = 31.22%; 95% CI = 4.01%). Based on the 95% CI REG pictures had more errors than EXC pictures, while there was no difference in error rate between REG and EXC words. The results of the mean error rate analyses indicated there were no significant speed-accuracy trade-offs.

#### Discussion

The results of Experiment 1 provide evidence that there is significant *shared* activation for words and pictures in the VWFA for EXC words and their corresponding pictures (Fig 4a), but not for REG words and pictures as the pictures were significantly more activated than the words while the words were not significantly activated in this region (i.e., *unique*; Fig 4b). However, there was some overlapping word and picture activation just posterior to this region, but it was greater (i.e., *dominant*) for pictures. In the LOC there was significant overlapping activation for both EXC pictures and their corresponding words, as well as REG pictures and words, but pictures produced significantly greater (i.e., *dominant*) activation than words in both cases (Fig 4c and 4d).

Overlapping activation in these regions supports the notion that words and pictures share a common functional architecture, and that the processes involved are overlapping in the ventral visual processing streams, while also supporting the dominance of processes involved in picture naming in the LOC compared to word reading. However, reading dominance was not established for words in the VWFA over picture naming, given that activation was significant but *shared* for EXC words and picture referents, and actually *unique* to pictures for REG stimuli. Furthermore, an additional analysis at the individual participant level found that no participants had significant activation for the Words > Pictures contrast within 10mm of the VWFA.

Finding any overlapping activation for words and pictures in these regions indicates that the VWFA and LOC reflect processes underlying both reading and picture naming to at least some degree. In the VWFA there is *shared* activation between words and pictures for EXC stimuli, but for REG stimuli pictures produce more activation than their corresponding words, which did not activate this region significantly, suggesting that EXC words are better suited for activating the VWFA than REG words, and that EXC word activation is more similar to picture activation than REG word activation is to picture activation in the VWFA. REG word activation of the VWFA may be less than EXC because sublexical (i.e., phonetic decoding) reading processes may sometimes be involved in REG word reading, resulting in less reliance on lexical processing. In the LOC, activation is greater for pictures than for words but words and pictures also have significant overlapping activation in the region (i.e., *dominant* picture activation).

In the behavioural analyses, REG pictures were named slower than EXC pictures, which is opposite to the typical regularity effect observed for words whereby REG words are read faster than EXC words. The error rate analysis also supported this effect, whereby REG pictures had more errors than EXC pictures. We suggest that this effect may reflect that representations for EXC words and their corresponding pictures enjoy a more direct connection between picture memory and the orthographic lexical system than REG words and pictures given that REG words may rely on dual processing routes (see Fig 1). Our lab is currently exploring a larger behavioural study, and this reverse regularity effect for pictures appears to be robust (Neudorf et al., in prep.).

Other research has demonstrated that physical differences between words and pictures (i.e., visual complexity) may introduce confounds in the evaluation of specialized mechanisms for the two classes of stimuli (Szwed et al., 2011). The use of stimuli that control for visual complexity was thus the focus of Experiment 2, to examine whether *dominant* activation remains in the LOC for picture naming under controlled visual complexity conditions.

### Experiment 2

Given that pictures have greater visual complexity than words (i.e., greater visual detail, broader range of spatial frequencies, and color), it may be the case that the differences in visual complexity between pictures and words could partly account for the results in Experiment 1 (Szwed et al., 2011). Due to the greater complexity in pictures, greater activation could occur with picture stimuli thus affecting any contrast with words to yield a larger value for pictures greater than words in any regions affected by visual complexity.

By superimposing words on the pictures and instructing the participant to name either the word or the picture, visual complexity was controlled for, as the same stimulus was used for both the reading and picture naming conditions (see Fig 3), providing a more stringent test of the contrast between pictures and words. Incongruent trials (50%) encouraged participants to inhibit processing of the stimulus type not being responded to in each block. Congruency effects were not modeled in the analysis because the number of trials did not produce enough power for an event related analysis, and we argue that the theoretically important element of the congruency manipulation was to ensure that participants were not relying on the other stimulus type while responding. However, the inclusion of these elements of attentional control and simultaneous stimulus presentation require additional cognitive processes not required in Experiment 1, which will be explored in our discussion of the results. Analogous to Experiment 1, Experiment 2 examined the overlap between processes underlying reading and picture naming using these superimposed stimuli. The hypotheses for Experiment 2 were identical to those in Experiment 1. However, if there are any differences in Experiment 1 due to visual complexity, then those will be equated in Experiment 2, thus increasing the precision of our comparison between word reading and picture naming.

#### FMRI Group Analyses

The factorial analysis of Word Type (EXC vs REG) x Stimulus Type (Pictures vs Words) produced a significant main effect for Pictures > Words only. The Pictures > Words contrast for EXC and REG stimuli separately did not identify any significant regions, so the factorial main effect was examined. The factorial Pictures > Words contrast was not significant in the VWFA, and pictures and words *shared* activation in this region (Fig 6a and 6b).

**Fig 6.**
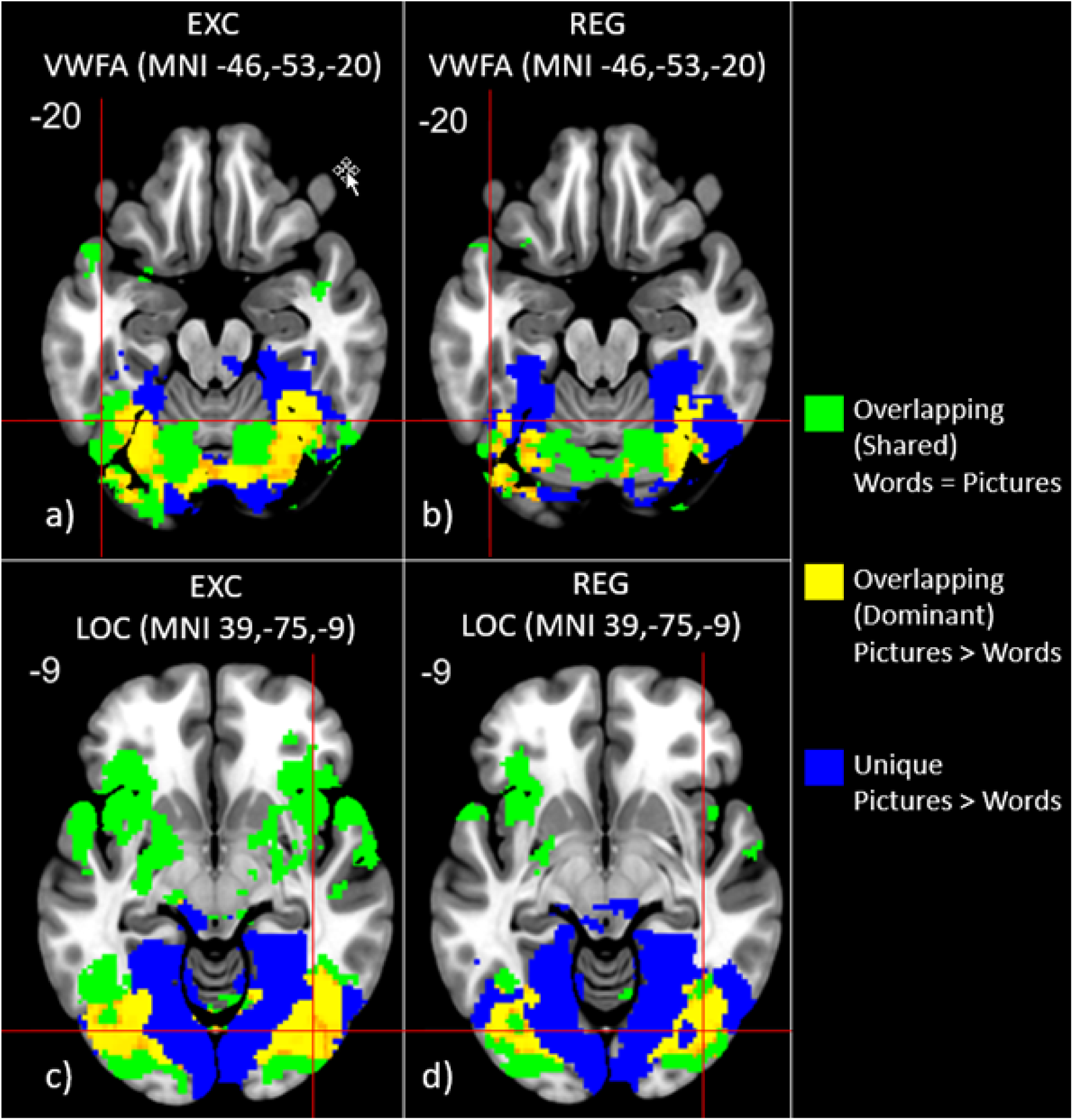
Experiment 1 Pictures > Words activation in VWFA (a,b) and LOC (c,d) for EXC (a,c) and REG (b,d) stimuli separately. Yellow represents an overlapping *dominant* effect for pictures (pictures minus rest) compared to words (words minus rest): both conditions were significant, but the effect for pictures was significantly higher than words. Blue indicated a *unique* effect for pictures (pictures minus rest) compared to words (words minus rest): the effect was only significant for pictures and a significant difference was detected between the effect for pictures and words. Green represents an overlapping *shared* effect for both pictures (pictures minus rest) and words (words minus rest): both conditions were significant, and the effect is not statistically different between pictures and words. There were no regions where Words > Pictures contrasts showed any dominant or unique activation. All images thresholded using TFCE with *p* < .01. *Dominant* activation values vary from dark orange (*p =* .01) to bright yellow (*p <* 0.001). *Unique* and *shared* activation values are binary based on whether the activation is significant at *p* < .01.

In the LOC, activation for pictures and words was primarily *shared*, but proximal regions (MNI: *x* = 33, *y* = -73, *z* = -14; and x = 43, y = -70, z = -7; both within 10mm of the center of LOC) demonstrated *dominant* picture activation (Fig 7).

**Fig 7.**
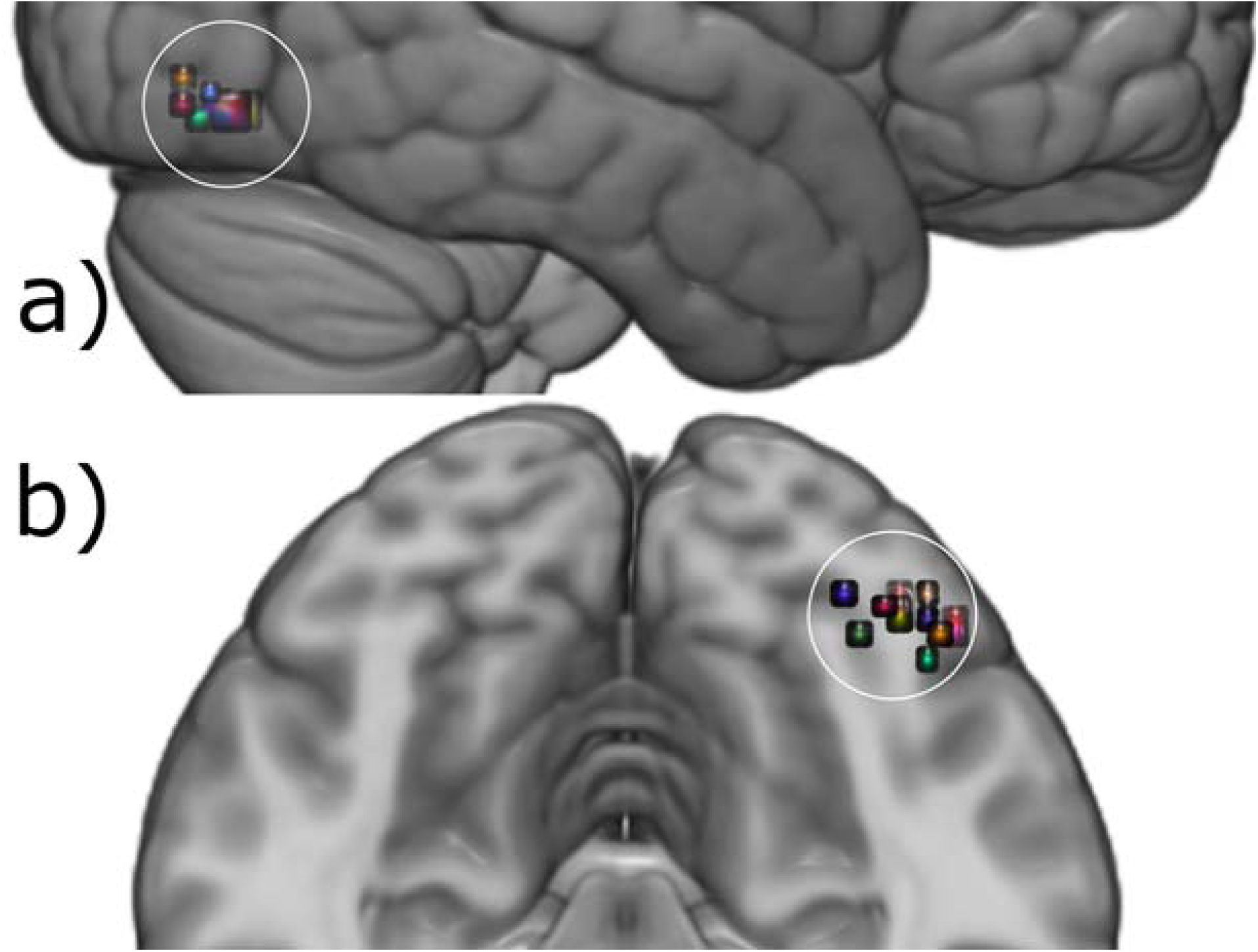
Experiment 1 sagittal (a) and axial (b) images of the individual localizer (Pictures > Words) ROI analysis of the LOC (10mm radius around MNI: x = 39, y = -75, z = -9). Fourteen out of 15 individuals showed activation in this region.

As in Experiment 1, other notable areas of *shared* word and picture activation for both EXC and REG words include activation of semantic regions in the anterior temporal lobes (see Fig 6 and Fig 7; see Patterson et al., 2007), activation of phonological regions in the inferior frontal gyri and insula (see Fig 6c, Fig 6d, and Fig 7; see Borowsky et al., 2006), and putamen activation in the basal ganglia (see Fig 6c, Fig 7c, and Fig 7d; see Gould et al., 2017, 2018; see also review by Price, 2012, and Oberhuber et al., 2013).

#### FMRI Individual Analyses

To investigate whether the Words > Pictures contrast may have been significant at the individual participant level in the VWFA, a ROI localizer analysis was conducted as in Experiment 1. Of the 15 participants, only 5 produced a significant Words > Pictures contrast within 10mm of the VWFA center, which is less than half and not significant from what would be expected by chance from the binomial distribution, which requires a threshold of 12 out of 15 to be met to demonstrate a consistent pattern of activation in the VWFA (see Fig 8 and Table 2). This finding is consistent with what was found at the group level, whereby no regions produced a significant Words > Pictures contrast in the VWFA.

**Fig 8.**
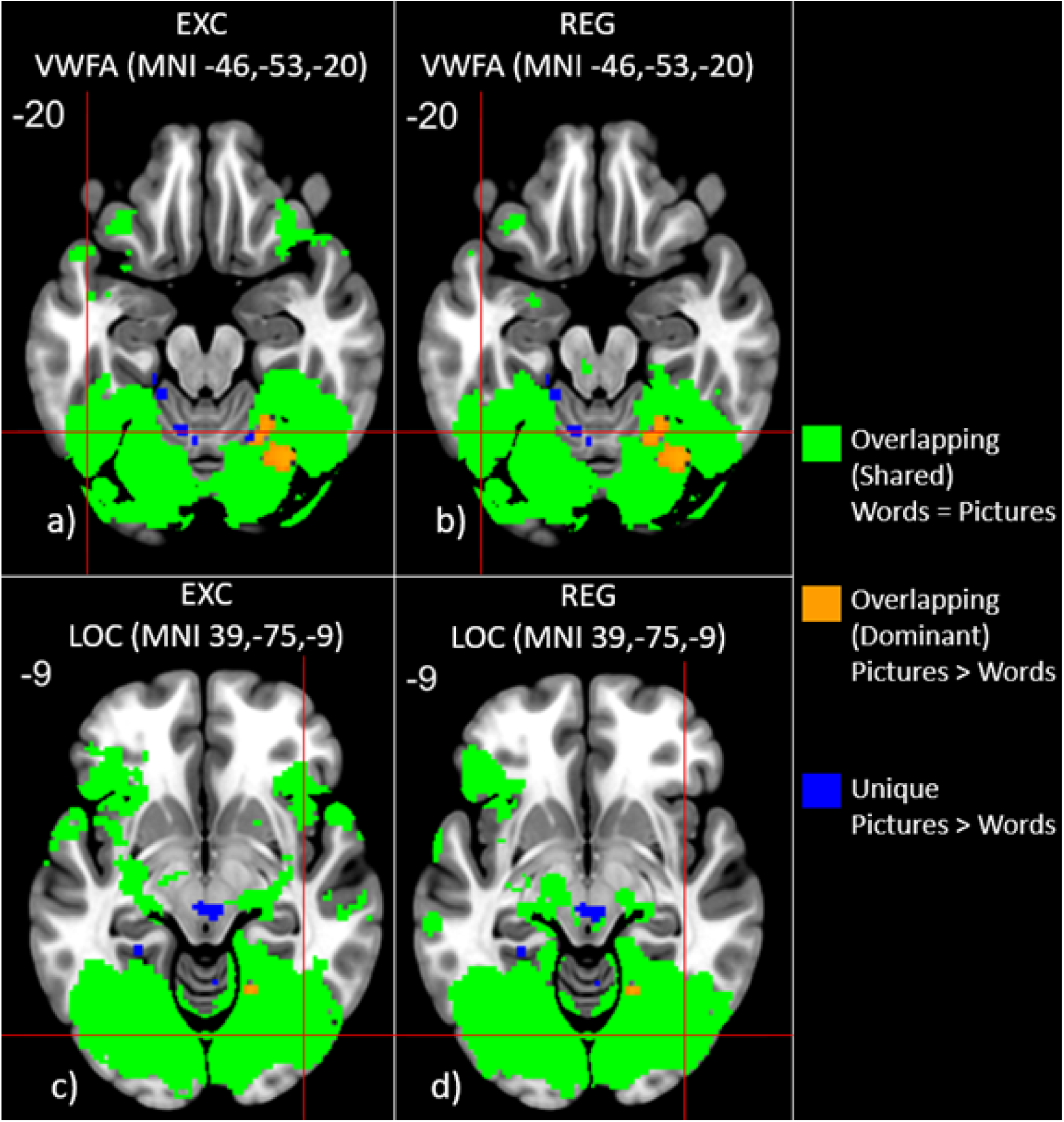
Experiment 2 factorial Pictures > Words activation in VWFA (a,b) and LOC (c,d) for EXC (a,c) and REG (b,d) stimuli separately. Orange represents an overlapping *dominant* effect for pictures (pictures minus rest) compared to words (words minus rest): both conditions were significant, but the effect for pictures was significantly higher than words. Blue indicated a *unique* effect for pictures (pictures minus rest) compared to words (words minus rest): the effect was only significant for pictures and a significant difference was detected between the effect for pictures and words. Green represents an overlapping *shared* effect for both pictures (pictures minus rest) and words (words minus rest): both conditions were significant, and the effect is not statistically different between pictures and words. There were no regions where Words > Pictures contrasts showed any dominant or unique activation. All images thresholded using TFCE with *p* < .01. *Dominant* activation values vary from dark orange (*p =* .01) to bright yellow (*p <* 0.001). *Unique* and *shared* activation values are binary based on whether the activation is significant at *p* < .01.

**Table 2.** Experiment 2 largest cluster coordinates (center of gravity; i.e. weighted average of the coordinates by activation intensity) of significant Word > Pictures activation within 10mm of VWFA (MNI: x = -46, y = -53, z = -20). Five of 15 participants had a significant contrast within a 10mm radius ROI centered on the VWFA.

| Participant | X | Y | Z | Voxels | Max Z-value |
| --- | --- | --- | --- | --- | --- |
| 2 | -51.6 | -53.8 | -23.9 | 50 | 5.70 |
| 5 | -45.5 | -58.8 | -26.0 | 5 | 6.23 |
| 6 | -42.5 | -47.5 | -21.0 | 4 | 4.10 |
| 12 | -45.6 | -60.7 | -17.4 | 15 | 5.36 |
| 13 | -44.6 | -54.6 | -25.7 | 6 | 4.82 |

Using the same approach for the Pictures > Words contrast in the LOC, 12 out of 15 participants produced a significant Pictures > Words contrast within 10mm of the LOC center, which meets the threshold of 12 out of 15 based on the binomial distribution (see Fig 9 and Table 3).

**Fig 9.**
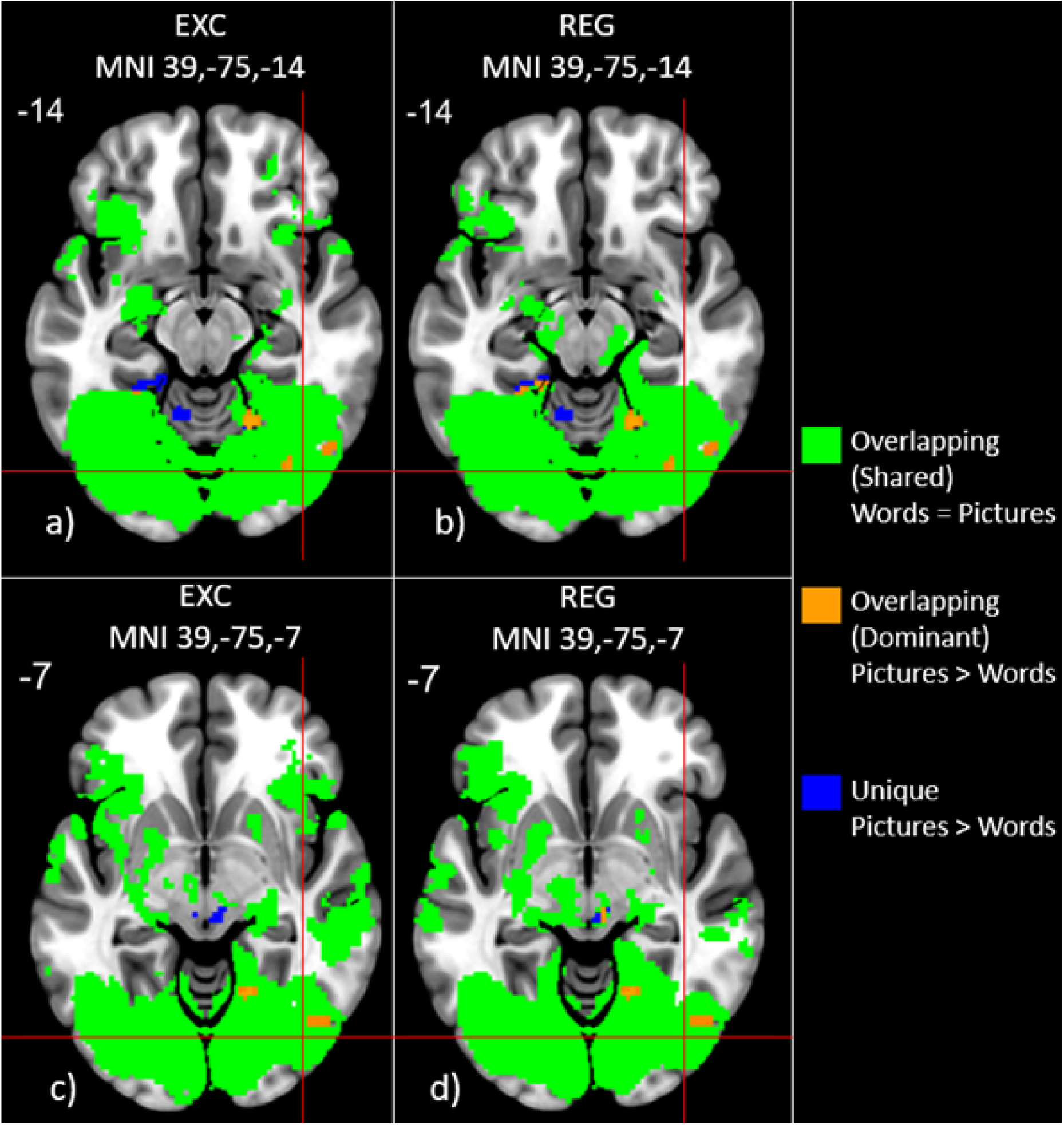
Experiment 2 factorial Pictures > Words activation just superior (a,b) and inferior (c,d) to LOC for EXC (a,c) and REG (b,d) stimuli separately. Orange represents an overlapping *dominant* effect for pictures (pictures minus rest) compared to words (words minus rest): both conditions were significant, but the effect for pictures was significantly higher than words. Blue indicated a *unique* effect for pictures (pictures minus rest) compared to words (words minus rest): the effect was only significant for pictures and a significant difference was detected between the effect for pictures and words. Green represents an overlapping *shared* effect for both pictures (pictures minus rest) and words (words minus rest): both conditions were significant, and the effect is not statistically different between pictures and words. There were no regions where Words > Pictures contrasts showed any dominant or unique activation. All images thresholded using TFCE with *p* < .01. *Dominant* activation values vary from dark orange (*p =* .01) to bright yellow (*p <* 0.001). *Unique* and *shared* activation values are binary based on whether the activation is significant at *p* < .01.

**Fig 10.**
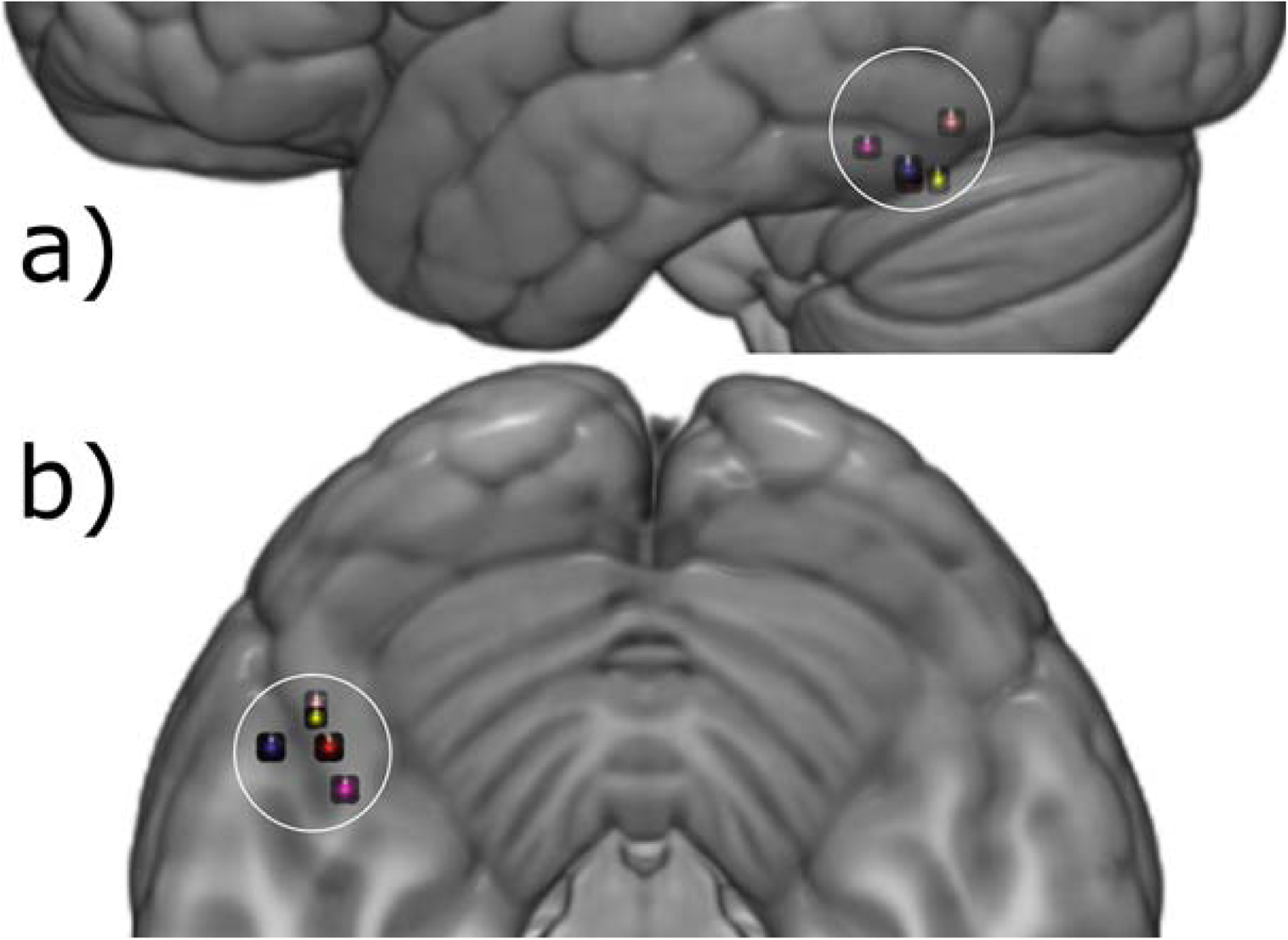
Experiment 2 sagittal (a) and axial (b) images of the individual localizer (Words>Pictures) ROI analysis of the VWFA (10mm radius around MNI: x = -46, y = - 53, z = -20). Only 5 out of 15 individuals showed activation in this region.

**Fig 11.**
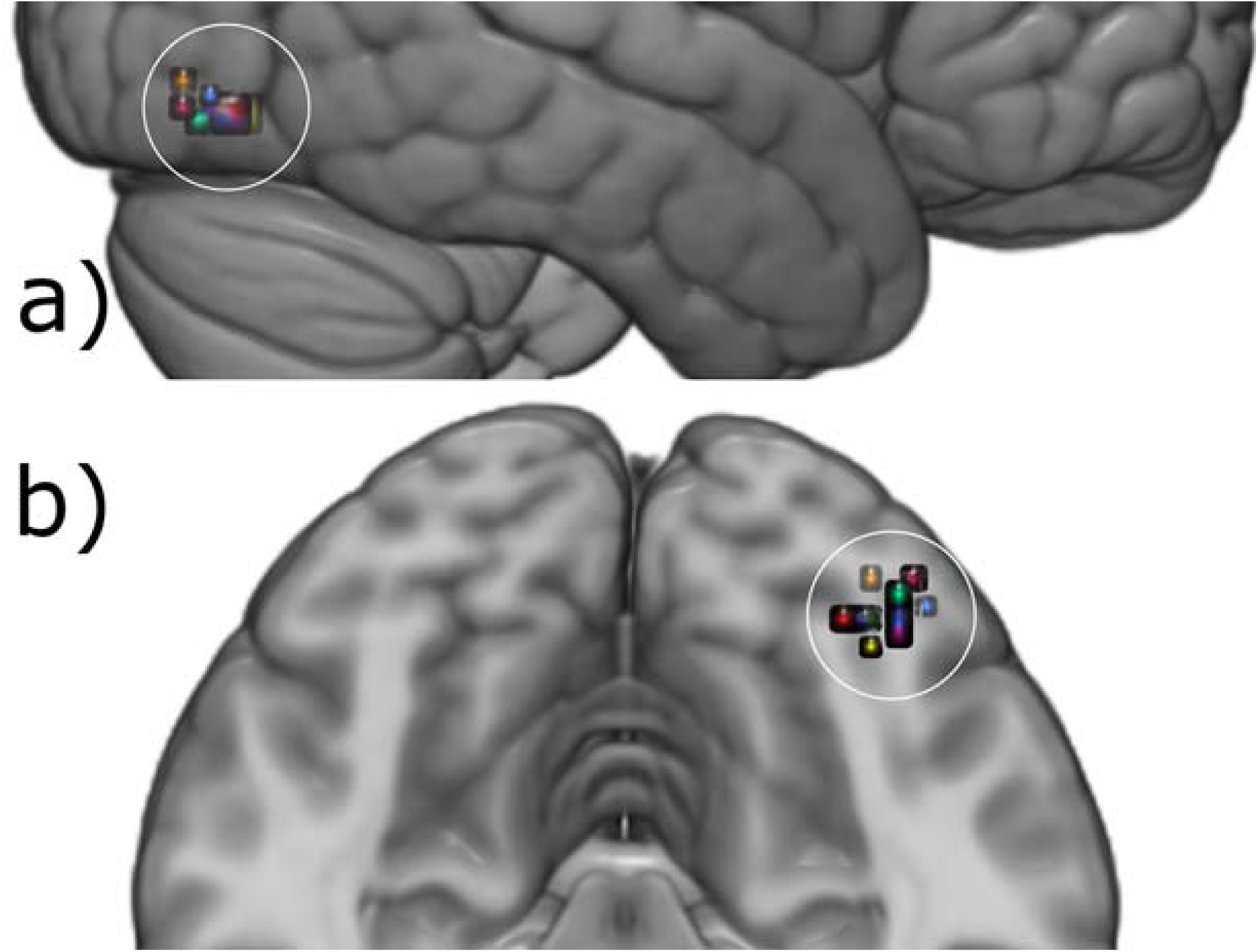
Experiment 2 sagittal (a) and axial (b) images of the individual localizer (Pictures > Words) ROI analysis of the LOC (10mm radius around MNI: x = 39, y = -75, z = -9). Eleven out of 15 individuals showed activation in this region.

**Table 3.** Experiment 2 largest cluster coordinates (center of gravity; i.e. weighted average of the coordinates by activation intensity) of significant Pictures > Words activation within 10mm of LOC (MNI: x = 39, y = -75, z = -9). Twelve of 15 participants had a significant contrast within a 10mm radius ROI centered on the LOC.

| Participant | X | Y | Z | Voxels | Max Z-value |
| --- | --- | --- | --- | --- | --- |
| 3 | 43.0 | -75.9 | -12.2 | 123 | 7.83 |
| 5 | 40.4 | -75.3 | -10.9 | 424 | 8.46 |
| 6 | 48.0 | -74.0 | -8.0 | 19 | 6.20 |
| 7 | 40.0 | -78.9 | -10.0 | 2 | 4.59 |
| 8 | 32.0 | -80.0 | -14.0 | 1 | 3.66 |
| 9 | 43.0 | -80.2 | -9.3 | 64 | 5.68 |
| 10 | 43.0 | -69.3 | -9.3 | 28 | 5.05 |
| 11 | 38.8 | -78.4 | -17.7 | 12 | 5.20 |
| 12 | 47.3 | -76.7 | -9.0 | 13 | 5.27 |
| 13 | 34.0 | -74.0 | -10.0 | 1 | 4.04 |
| 14 | 46.0 | -74.0 | -16.0 | 1 | 3.67 |
| 15 | 40.4 | -75.3 | -10.9 | 424 | 8.46 |

#### Reaction Time

Nine participants’ behavioral data were unaffected by the faulty connector described earlier, so their data was analyzed with the caution noted that the sample size is small. There was a significant main effect of Stimulus Type, *F*(1, 7) = 35.6, *MSe* = 6793.2, *p =* .001, whereby words (*M* = 729.18 ms) are named faster than pictures (*M* = 903.65 ms), which is consistent with Experiment 1 and previous reports in the literature. There was no significant main effect of Word Type, *F*(1, 7) = 3.16, *MSe* = 2302.84, *p* = .12, and no significant Stimulus Type x Word Type interaction, *F*(1, 7) = 2.19, *MSe* = 2033.74, *p* = .18 (REG Words, *M* = 702.26 ms; EXC Words, *M* = 756.10 ms; REG Pictures, *M* = 900.52 ms; EXC Pictures, *M* = 906.79 ms; 95% CI = 25.15 ms). Based on the 95% CI, there was no significant difference between EXC and REG pictures, but REG words were read significantly faster than EXC words, as is typically reported in the literature (see Cummine et al., 2010 for an example and a review).

#### Error Rate

There was a significant main effect of Stimulus Type, *F*(1, 7) = 8.10, *MSe* = 84.5, *p* = .02, whereby there were fewer errors in the word reading conditions (*M* = 3.00%) than the picture naming conditions (*M* = 12.23%). There was no significant main effect of Word Type, *F*(1, 7) = .46, *MSe* = 87.36, *p* = .52, and no significant Stimulus Type x Word Type interaction, *F*(1, 7) =.96, *MSe* = 87.93, *p* = .36 (REG Words, *M* = 2.52%; EXC Words, *M* = 3.48%; REG Pictures, *M* = 14.97%; EXC Pictures, *M* = 9.49%; 95% CI = 5.23%). Based on the 95% CI, REG pictures had more errors than EXC pictures, but there was no significant difference between REG and EXC words. The results of the mean error rate analyses indicated there were no significant speed-accuracy trade-offs.

#### Discussion

Experiment 2 provides further evidence that there is significant *shared* activation for words and pictures in the VWFA for EXC words and their corresponding pictures, and also for REG words and pictures. In the LOC there was predominantly *shared* activation for both EXC and REG pictures and their corresponding words, but pictures produced regions with significantly greater (i.e., *dominant*) activation than words in the vicinity of LOC.

Shared patterns of activation could be expected given that the stimuli were matched for both conditions, and that the phonological output was as similar as possible between conditions. However, participants were encouraged to ignore the other stimulus type due to the non-predictive (50%) congruency. Even so, the dominance of picture processing was demonstrated in the LOC compared to word processing. Picture dominant processing was also found in more medial regions of the fusiform gyrus, supported by research demonstrating silent naming of inanimate object pictures elicited greater activation than animal pictures (Martin & Chao, 2001). Evidence from the stroke literature suggests that many distributed regions of the temporal cortex are involved in picture naming (DeLeon et al., 2007), which is consistent with our findings of Picture dominant regions not just near the right LOC but in bilateral regions of the temporal lobe for both experiments. However, word dominance was not established for words in the VWFA over picture naming, given that activation was significant but *shared* for words and picture referents. There was no other evidence for either *dominant* or *unique* activation in the VWFA or the LOC.

Furthermore, an additional analysis at the individual participant level found that only 5 participants had significant activation for the Words > Pictures contrast within 10mm of the VWFA, which did not meet the threshold of 12 out of 15 participants from the binomial distribution. Looking at the LOC, the individual analysis found that 12 participants had significant activation for the Pictures > Words contrast within 10mm of the LOC, which meets the threshold of 12 out of 15 participants.

In the behavioural analyses, pictures were named slower and had more errors than words. The typical regularity effect was observed whereby REG words were read faster than EXC words, but no difference was observed between REG and EXC RT for pictures as was found in Experiment 1, but this null effect could be due to the small sample size. However, REG pictures elicited more errors than EXC pictures, as was observed in Experiment 1. The finding that picture-orthography agreement was unrelated to Word Type confirms that in Experiment 1 the slower RT for REG pictures than EXC pictures and that in Experiments 1 and 2 the greater number of errors for REG than EXC pictures was not related to the agreement between the pictures and words used. It may be that the connections between picture memory and the orthographic lexical system are inhibited in this combined presentation version of the task given the 50% incongruity between pictures and words, but future research should explore this possibility further through manipulation of the congruity proportion.

The *dominant* activation found in the LOC in Experiment 2 was for pictures, which supports the notion that at least part of the LOC region is *dominant* for picture processing (e.g., Grill-Spector et al., 2001; Kanwisher et al., 1996; Kanwisher & Dilks, 2013; Malach et al., 1995). Taken together, data from Experiment 1 predominantly supports the *shared* reading and picture naming hypotheses for both VWFA and for LOC, while demonstrating a greater sensitivity to pictures than words in regions of the LOC, and Experiment 2 extends and replicates Experiment 1 and provides converging evidence for greater picture than word activation in the LOC. To the extent that superimposed word-picture stimuli controlled for visual complexity, Experiment 2 provided strong converging evidence. However, it is important to note that any extra shared activation in Experiment 2 relative to Experiment 1 could also be due, at least in part, to the presence of both a word and a picture on each trial.

## General Discussion

These experiments examined how lexical reading along the left VWFA has either *unique* or overlapping (*shared* or *dominant*) activation loci relative to picture naming of the same referents, and how picture naming along the right LOC has either *unique* or overlapping activation loci relative to word versions of the same referents. *Unique* activation describes when one condition produces significant activation while the other does not, and there is a statistically significant contrast between the two conditions. Overlapping activation describes when both conditions produce significant activation, and this is described as *shared* when there is no significant contrast between the conditions, and *dominant* when there is a significant contrast in favor of one of the conditions.

When stimuli were presented individually in Experiment 1, the group analysis showed that the VWFA was activated in a *shared* fashion for EXC words and pictures. Given the common characterization of the VWFA to be more sensitive to word stimuli, and given that EXC words represent the optimal “lexical” stimuli, this finding calls this common characterization into question. The use of matched EXC words and pictures provides a valuable addition to research seeking to understand lexical word versus picture processing in the ventral stream. These results suggest that the VWFA is not as specialized for words compared to pictures as previously proposed. The LOC was activated in a predominantly *dominant* fashion by pictures when the stimuli were presented in isolation. The LOC and VWFA were activated in a predominantly *shared* fashion by words and their corresponding pictures when they were superimposed in Experiment 2 (although this could be expected more so than in Experiment 1, as both words and pictures were being presented on each trial), with the primary result being the region of picture *dominant* activation, which provides converging evidence for what was seen in Experiment 1. Given that activation patterns changed across experiments, it suggests that visual complexity does have an impact on these effects in the LOC and VWFA, and in particular there is not as much *dominant* activation for pictures over words in the LOC when visual complexity is controlled. In the individual VWFA analyses, there were no participants that had a region of activation where words had greater activation than pictures in Experiment 1 (0 out of 15 participants) and a non-significant proportion in Experiment 2 (5 out of 15 participants), consistent with the lack of VWFA word *dominant* or *unique* activation in the group analyses. In the individual LOC analyses, a significant number of participants had a region of activation where pictures had greater activation than words in Experiment 1 (14 out of 15), and in Experiment 2 (12 out of 15). Taken together, these results support the notion that the LOC is involved more so in picture naming, but is also sensitive to word form.

The general finding of overlapping functional processing between words and pictures is consistent with the variety of patient data in which one can have visual object agnosia with preservation of reading abilities (Gomori & Hawryluk, 1984), or show impairments in both modalities (e.g., patients with pure alexia are often reported to have difficulties with color naming and picture processing; Behrmann et al., 1998; Damasio & Damasio, 1983; De Renzi et al., 1987; Geschwind, 1965), or selective impairment in reading while having spared picture, color, and letter naming (Benson & Geschwind, 1969; although Friedman & Alexander, 1984 found that picture identification was impaired at faster stimulus presentation speeds, suggesting that visual identification deficits may not always be apparent at long stimulus presentation times; see also Price et al., 2006 for a discussion of how patient data allows for the assessment of *necessity* of a brain region, whereas functional imaging data in healthy participants can only assess *sufficiency* of a brain region). Given that the patient data suggests that reading deficits and picture naming deficits are mathematically independent (i.e., one can have either deficits in either picture processing, reading, or both), it would seem that the overlapping activation found in our research is consistent with the existence of all three patient types if we consider the overlapping activation to reflect some degree of redundancy in reading and picture naming systems. The reading system has likely co-opted preexisting regions that originally subserved object or picture identification and naming (i.e., “the neuronal recycling hypothesis”, e.g., Dehaene & Cohen, 2011), which may be responsible for some degree of redundancy between word reading and picture naming.

Presenting stimuli both in isolation and superimposed is a useful manipulation in order to equate the influence of visual complexity. One advantage of presenting these stimuli superimposed was that it allowed us to determine whether *shared* activation could be accounted for simply by superimposing stimuli. While this may partially account for some of the *shared* activation within the LOC for superimposed stimuli, it is important to note that there was still significant overlapping activation in the LOC when words and pictures were presented in isolation, although picture activation was *dominant*. In contrast, the VWFA showed a similar pattern of results regardless of whether the stimuli were presented in isolation or superimposed for the EXC stimuli, while the REG stimuli demonstrated a lack of word activation in the isolated presentation and a pattern of *shared* activation with the superimposed presentation, suggesting that REG words are not ideal for activating the VWFA region. This observation may be important for interpreting past research showing greater activation in VWFA for pictures than words, given that REG versus EXC word types were not manipulated (e.g., Price & Devlin, 2003). However, note that the EXC words did not produce a statistically significant contrast compared to REG words, likely due to them both having relatively high frequencies of occurrence in print, so further research employing a wider range of frequency of occurrence needs to be done to determine whether this is a robust effect in the VWFA. However, there is a theoretical rationale to rely more on EXC words given that they must be read lexically to be read correctly, whereas REG words can be read correctly by sublexical processing. Nonetheless, it has proved important to include both isolated and superimposed stimulus presentation conditions in order to evaluate the influence of these factors.

Experiment 2 is effectively the same as the picture-word interference paradigm (on which there is a large body of research; e.g., (Costa et al., 2005; Cummine et al., 2013; G. de Zubicaray et al., 2002; Gauvin et al., 2018). Neuroimaging research of the picture-word interference paradigm, whereby related or unrelated words are superimposed on a picture to be named, has demonstrated a word frequency effect in bilateral premotor, primary sensorimotor, anterior cingulate cortex, left posterior superior and middle temporal gyrus, supplementary motor area, and posterior temporal cortex, and right supramarginal gyrus and posterior superior temporal gyrus (de Zubicaray et al., 2012). These are primarily dorsal regions, with the most ventral being the left middle temporal gyrus, whereas our VWFA and LOC ROIs are more ventrally located in the inferior temporal gyrus. Considering that we have analyzed both the congruent and incongruent stimuli together, we are not explicitly measuring the effects of semantic relatedness between word and picture stimuli, but it is important to consider how the paradigm influences the processes involved in responding to the stimuli. In addition to our primary goal of matching the stimulus complexity, it could be argued that this paradigm also requires participants to inhibit processing of the stimulus which they are instructed not to respond to. Additionally, by viewing both stimulus types simultaneously, both pathways may be jointly stimulated. Under the assumption of stimulus inhibition, responses to words should have a better chance to show more activation in the VWFA than pictures to the extent that the participant is able to effectively inhibit the picture modality. In the case that both pathways are jointly stimulated, there should be shared activation at a large scale. In general we found much more shared activation in Experiment 2 than in Experiment 1, but in Experiment 1 the VWFA had prominent picture dominant and unique regions of activation, so the large amount of shared activation in the VWFA for Experiment 2 represents a better representation of word activation with stimuli combined than when stimuli were presented separately. Additionally, in the individual analyses we found that in Experiment 1 there were no individuals with words greater than pictures contrasts in the VWFA, but in Experiment 2 there were 5 out of 15 individuals showing this contrast, representing a better opportunity for this contrast to be observed. These observations in the VWFA seem to support that by controlling stimulus complexity and creating a paradigm which may have resulted in participants inhibiting the other stimulus, the opportunity was improved for the words greater than pictures contrast, but ultimately it was not observed. Even with the more level playing field this paradigm produced, pictures still produced greater activation than words in regions of the LOC. It also clear, however, that both the VWFA and LOC are being recruited in a more shared fashion when reading words and naming picture stimuli, suggesting that both word reading and picture naming streams of processing are being activated to some extent during this paradigm.

Conjunction analyses of fMRI data are an important strategy employed by many researchers (e.g, see Price & Friston, 1997 seminal work on conjunction analyses; see also Borowsky et al., 2005 for another variant). The technique used here not only delineates between *unique* and overlapping activation, but also specifies whether the overlapping activation was *dominant* for one condition or not (*shared*), and was helpful in identifying where activation was more sensitive to picture naming than to word reading in the LOC, even though activation was overlapping (*dominant*). This approach may be useful for other researchers in the future who wish to describe the nature of a contrast (*unique* or *dominant*), and to describe the nature of a binary conjunction (*shared* or *dominant*). These concrete definitions allow for a useful hybrid approach to contrast analyses and conjunction analyses.

A localizer approach to studying regions such as the VWFA and LOC has been widely used, but verification of the important underlying assumption that individual localized regions are consistent enough to be labeled as the same region has not. By using the approach used in these experiments to verify that a significant number of individuals show consistent activation within a reasonable distance from the expected location of the ROI, the validity of such a localizer approach can be readily assessed.

In assessing *shared* processing in these regions of interest, it is important to note that both the VWFA and LOC are often considered to be well beyond low-level visual feature processing systems, and well before high-level (semantic) language processing. Rather, the VWFA has been argued to specialize in word form identification and abstract representations of visual words (e.g., Cohen et al., 2000; Dehaene et al., 2001, 2002; Dehaene & Cohen, 2011), and the LOC has been argued to be an object-selective region, as it responds more strongly to pictures of objects than to their scrambled counterparts, appears to be object-invariant, and shows a number of responsive properties that characterize an effective object recognition system (e.g., Grill-Spector et al., 1999; Kourtzi, 2001; Malach et al., 1995; Snow et al., 2011). Based on these arguments, the *shared* activation found in these regions would not likely be attributable to either low-level feature processing or high-level semantic processing. The overlapping activation in these regions for word reading and picture naming processes are revealing in terms of how these processes operate in the brain, and suggests that they may not be as specialized as previous research, and models, have suggested.

### Limitations

It is also important to consider that greater activation for one task than another task does not always mean that region is more sensitive to the one task, but may alternatively mean that processing time is longer for that task or more effort is required for that task (e.g., see Taylor et al., 2013 for a discussion of these effects of engagement). For example, as has been demonstrated in past research (e.g., Fraisse, 1969; Hennessey & Kirsner, 1999; Potter & Faulconer, 1975) and as was reported in our reaction time results for Experiments 1 and 2, words were named faster than pictures, which suggests more processing time and/or effort is required for naming pictures.

Additionally, there may have been differences in semantic processing, given that word reading does not necessarily involve the same amount of semantic processing as picture naming because word reading can rely on direct orthography to phonology mapping. However, depending on how prototypical a picture is, it may be represented symbolically and thus named automatically without relying on semantics (e.g., naming a simple picture like “%” as “percent” without requiring the semantic activation of the concept of ratio portrayed its image). Other imaging modalities such as magnetoencephalography (MEG) with better temporal resolution than fMRI and better spatial resolution than electroencephalography (EEG) have demonstrated the ability to use a time-resolved approach to identify different stages of processing in specific locations including semantic processing of verbal sentences starting in the left anterior temporal lobe, then moving more posterior in the temporal lobe, before propagating to the left inferior frontal gyrus (e.g., Leonardelli et al., 2019; Lyu et al., 2019). These approaches could be used to address questions about which stages of processing elicit differential activation between word reading and picture naming in regions of the ventral stream such as the VWFA and LOC.

Using univariate methods, the current study is able to investigate brain activity levels but not necessarily information content in various brain regions. Other research has observed conceptual (i.e., semantic) overlap in visual word and picture processing (e.g., Devereux et al., 2013; Fairhall & Caramazza, 2013) using multivariate approaches such as multi-voxel pattern analysis (MVPA). This multivariate approach has also been able to show ventral occipital discriminability between words and false-fonts where univariate methods could not (Nestor et al., 2013). Based on the limitations noted whereby RT differs between picture and word conditions, and that size and visual complexity of pictures likely elicits more cortical activity than words in general, studies designed to take advantage of multivariate analysis approaches would be beneficial for confirming whether there are differences in word versus picture information in the fusiform gyrus irrespective of the overall amount of activity.

### Future Directions

Behavioural measurements of RT and error rates in both experiments demonstrated that RT (Experiment 1) and error rate (Experiments 1 and 2) show poorer performance for REG pictures than EXC pictures, which is opposite to the typical regularity effect for words whereby REG words are read faster than EXC words, as was observed in Experiment 2. This effect for pictures may be due to more optimal connections between picture memory and the orthographic lexical system for EXC words than for REG words, given that EXC words rely on the ventral-lexical stream, whereas REG words can rely on both or either of the ventral-lexical and the dorsal-sublexical streams (see Fig 1). This theory should be investigated further in future research, and our lab is currently exploring a larger behavioural study that investigates this issue.

Several studies have examined the conceptual (i.e., semantic) overlap in visual word and picture related tasks (Bright et al., 2004; Devereux et al., 2013; Fairhall & Caramazza, 2013; Shinkareva et al., 2011; Vandenberghe et al., 1996). For example, Bright et al. (2004) used PET to examine whether separate conceptual representations exist between words and pictures, or whether inputs from the different modalities converge on the same set of representations. The results demonstrated robust activation common to both modalities in anterior and medial aspects of the left fusiform gyrus, left parahippocampal cortex, left inferior frontal gyrus, and the anterior temporal lobes. These data are consistent with our data as we also found *shared* word and picture activation in the fusiform gyrus, inferior frontal gyrus, and anterior temporal lobe, and support the notion of a unitary system of semantic representations for both words and pictures, beginning in the occipital lobe and processed along the ventral stream and forward to the anterior regions of the inferior temporal cortex. The idea of a gradient in the ventral stream from perceptual sensitivity to categorical or semantic sensitivity has been supported by both word reading (Borghesani et al., 2016) and semantic question tasks (Martin et al., 2018), which have shown that visual perceptual differentiation occurs more in more posterior regions while more categorical distinctions occur in more anterior regions as processing moves towards the anterior temporal lobe. Borghesani et al. (2016) demonstrated that although processing becomes more abstract and category sensitive in anterior regions, regions from the occipital-temporal cortex all the way back to the primary visual area of the occipital lobe are sensitive to semantic category distinction between animals and tools. Martin et al. (2018) also found a shift from specific visual semantic distinction in the LOC to broad categorical semantic sensitivity in the anterior temporal lobe and demonstrated that visual features of object referents of words were represented in the LOC during a semantic property verification task. Given that semantic processing has been shown to occur at some level all the way from the occipital lobe to the anterior temporal lobe along the ventral stream, our finding that the processes involved in word reading and picture naming activate overlapping regions within the VWFA and LOC in Experiments 1 and 2, is generally consistent with this research. Future research that compares novel words paired with novel picture stimuli, while manipulating the degree of semantic representation (i.e., a learning paradigm whereby some novel words and pictures are given a minimal amount of semantic attributes, while others are given a greater amount), may be ideal for disentangling the effects of overlap due to the posterior-anterior semantic gradient along the ventral stream, from pure word form and picture form effects. The observed overlapping activation in the cortex for reading and picture naming may also reflect the co-opting of picture identification and naming for the purpose of reading (i.e., picture imagery) as well as the co-opting of visual word form identification and reading for the purpose of picture identification (e.g., word form imagery). Future research including word and picture imagery conditions would be important for assessing this possibility, given that mental imagery produces activation similar to visual perception (e.g., Ganis et al., 2004; Kosslyn et al., 1993; see also Mattheiss et al., 2018 for a method of localizing regions of visual semantic processing using ratings of word imageability).

## Conclusion

This research makes important contributions to fMRI research techniques, including the definition and demonstration of using *unique, dominant,* and *shared* activation maps to portray a combination of information about contrasts and conjunctions. Additionally, an important test was demonstrated of the underlying assumption of some localizer approaches that the individual localized regions are located within a constrained, consistent region. The present experiments provide new information regarding the overlap between processes underlying word reading and picture naming. Overall, the *shared* activation for words and pictures in the VWFA for group and individual analyses suggests word reading may share a common functional architecture more so than current models suggest (see Nestor et al., 2013 for similar findings regarding the overlap of representations for words and faces in the VWFA, with no specialized regions observed). Converging evidence from group and individual analyses of the LOC suggested consistent picture *dominant* activation near the LOC center and in medial regions of the fusiform gyri, even in Experiment 2 when visual stimuli were equated between conditions. This research also underscores the need for future research to include EXC word and picture stimuli for comparing lexical-based reading to picture naming, as REG words may be less than optimal given that they can also be read through the dorsal sublexical pathway. Given that we regularly engage in both word reading and picture naming together in our day-to-day experiences, and that children often learn to read with the aid of matching words and pictures, the ecological validity of studying them together is also an imperative.

## Acknowledgments

This work was supported by the Natural Sciences and Engineering Research Council (NSERC; http://www.nserc-crsng.gc.ca) of Canada in the form of NSERC Canada Graduate Scholarships to J. Neudorf, L. Gould and C. Ekstrand, and an NSERC Discovery Grant to the senior author, R. Borowsky under Grant 183968-2013-22. M. Mickleborough was also supported through a Post-Doctoral Research Fellowship from the Saskatchewan Health Research Foundation (SHRF). The lead author (J. Neudorf) and second author (L. Gould) contributed equally to this work. Correspondence can be sent to. The funders had no role in study design, data collection and analysis, decision to publish, or preparation of the manuscript.

## Supporting information

**S1 Table.** Regular and exception word stimuli.

| Regular Words | Exception Words |
| --- | --- |
| brain | bear |
| cliff | blood |
| coin | bowl |
| couch | bread |
| crib | bull |
| cube | comb |
| flame | door |
| flea | foot |
| girl | glove |
| hand | geese |
| harp | heart |
| hoop | hook |
| leaf | mould |
| match | pear |
| mouth | pint |
| mug | shoe |
| plum | soup |
| pope | sponge |
| pork | steak |
| shed | thread |
| slug | wasp |
| stump | wolf |
| toad | wood |
| toast | wool |
| tooth | world |

**S2 Fig.**
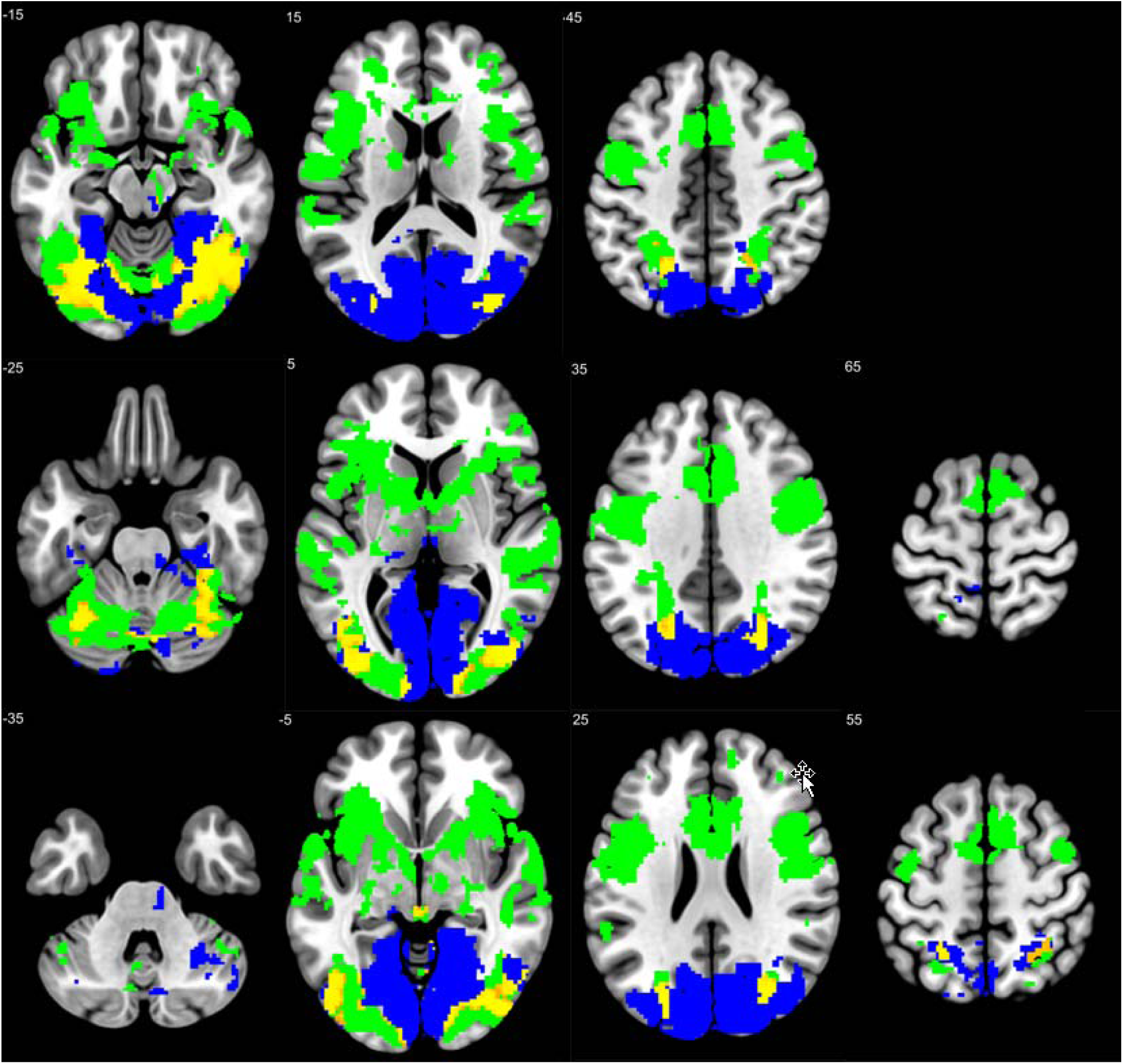
Experiment 1 EXC additional axial slices.

**S3 Fig.**
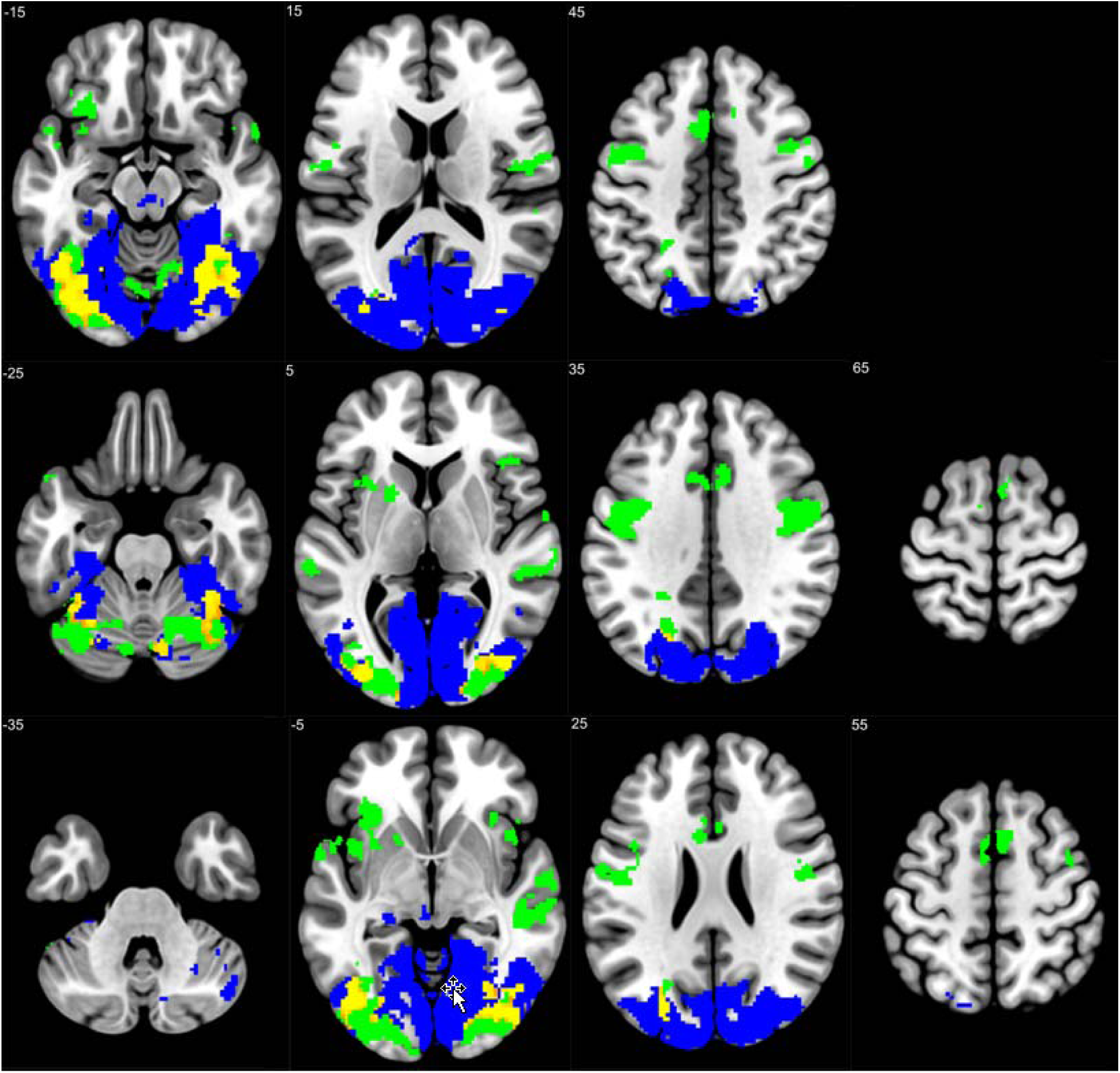
Experiment 1 REG additional axial slices.

**S4 Fig.**
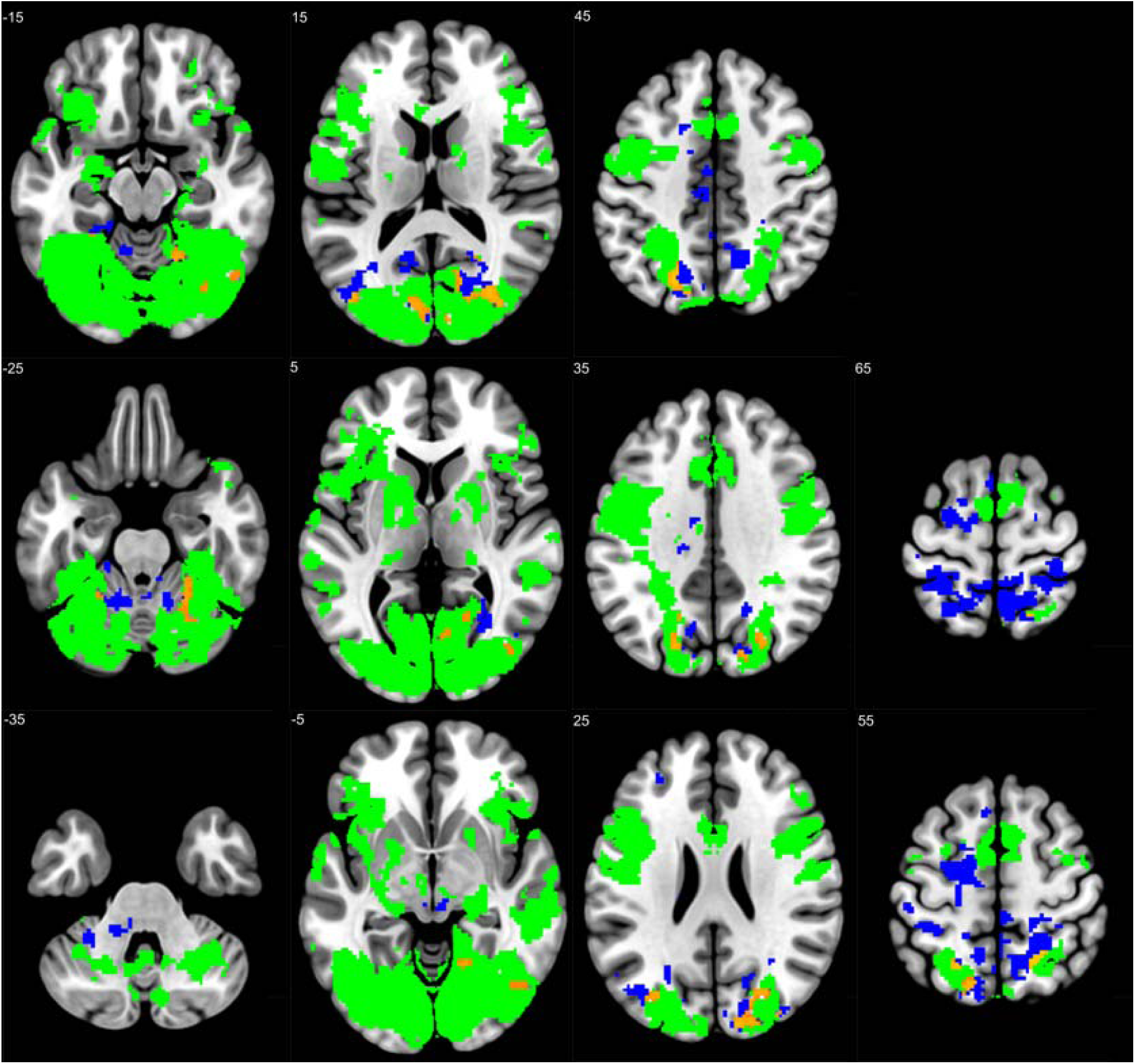
Experiment 2 EXC additional axial slices.

**S5 Fig.**
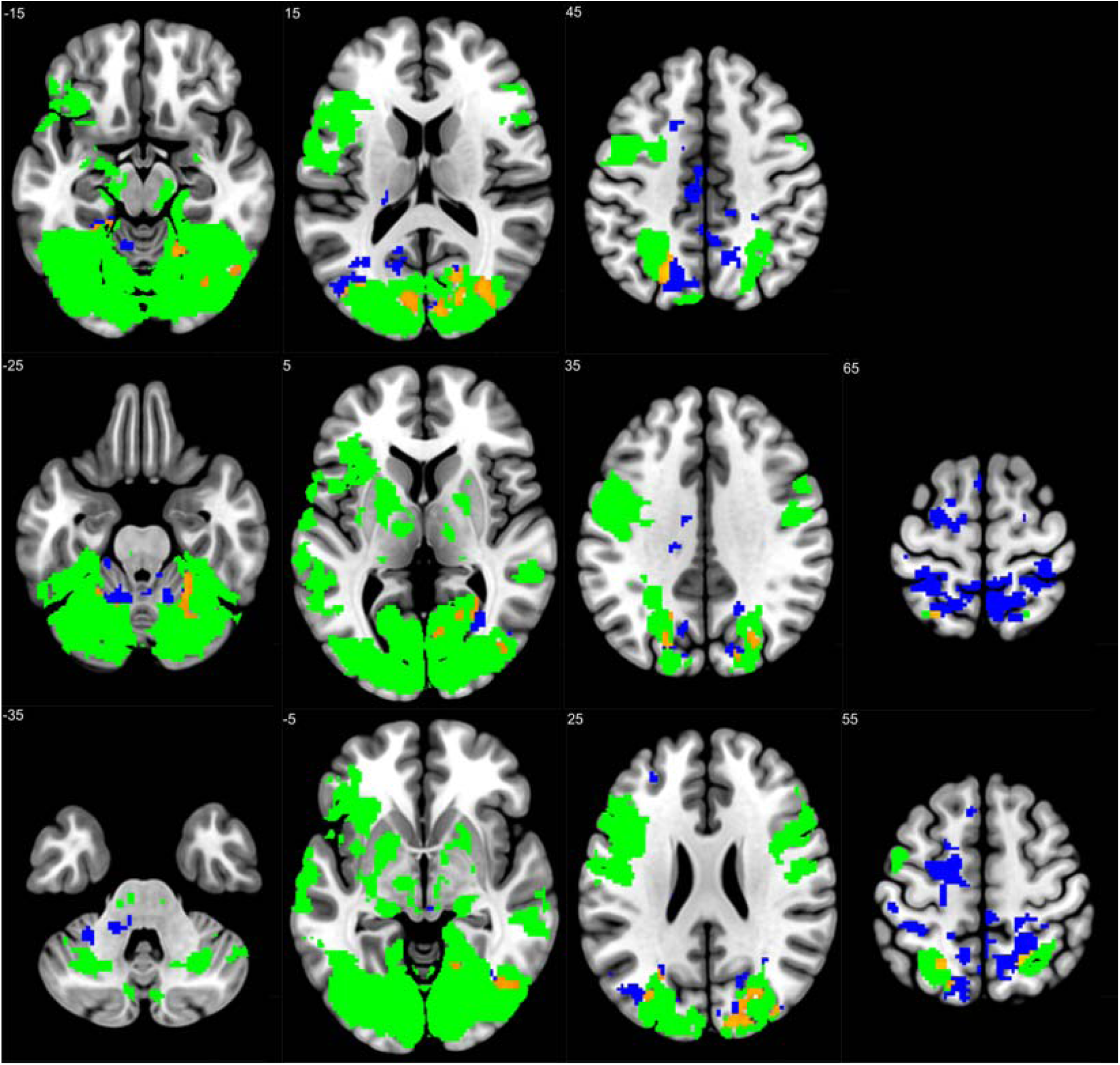
Experiment 2 REG additional axial slices.

